# Single-Nucleus Transcriptomics of the Mouse Medial Preoptic Area Reveals Sex-Dependent Molecular Signatures of Social Dominance

**DOI:** 10.64898/2026.04.30.721920

**Authors:** Tyler M. Milewski, Kathryn M. Mahach, Isaac Miller-Crews, Hans A. Hofmann, James P. Curley

**Author notes:** These authors contributed equally to this work (Tyler M. Milewski, Kathryn M. Mahach, Isaac Miller-Crews).

## Abstract

Social dominance hierarchies are a common form of social organization across animal species. We have previously shown that both male and female CD-1 mice form highly linear dominance hierarchies. In mice and other vertebrates, the medial preoptic area (mPOA) is a key hypothalamic sub-region regulating aggressive and defensive behaviors that support hierarchical social structures, but the transcriptional mechanisms in mPOA neurons underlying dominance behaviors and the influence of sex on these neuron populations in the context of social dominance hierarchies remain largely unresolved. Using single-nucleus RNA sequencing (snRNA-seq) to profile mPOA neurons from dominant and subordinate mice, we identified highly consistent social status-dependent changes in the transcriptomes of neuronal nuclei expressing neuropeptide transcripts. Oxytocin expression was remarkably widespread across mPOA neurons, and we found it to be the primary driver of group differences in neuropeptide co-expression networks. Overall, dominant males and females exhibited markedly decreased expression of oxytocin and vasopressin and had a lower proportion of neurons co-expressing multiple neuropeptide transcripts compared to subordinate individuals. In contrast, subordinates displayed widespread reorganization of the transcriptomic neuropeptidome and strikingly enhanced coupling of neuropeptide expression in mPOA neurons. Despite the strong pattern of concordant gene expression in dominant and subordinate individuals, the number of genes that were differentially expressed by status was substantially reduced in males compared to females. In sum, these results demonstrate that maintenance of social status dynamically reconfigures hypothalamic transcriptomic neuropeptidome in a sex-dependent manner and establishes how social status is encoded at a single-cell resolution.

## INTRODUCTION

Ranking systems, such as social dominance hierarchies, are common across animal taxa as they allow members of social groups to compete for access to resources essential for fitness in a predictable and organized manner. As dominant-subordinate relationships form and individuals taken on different social status roles, dominant and subordinate animals undergo coordinated and sex-specific shifts in physiology and behavior (Huffman et al., 2012; Maruska, 2015; Milewski et al., 2022; Morin et al., 2024). For example, dominant individuals often increase territorial patrolling, aggression, urination, scent-marking, feeding and drinking, whereas subordinates may shift toward avoidance and altered space use, adjust metabolic rates, feeding and drinking schedules, and show changes in sleep-wake patterns and stress-related physiology (Milewski et al., 2022).

The medial preoptic area (mPOA) of the anterior hypothalamus is an evolutionarily conserved brain region responsible for the coordination of social behavior including scent marking and aggression and contributes to the regulation of homeostatic processes such as feeding, drinking, sleep, stress regulation, and metabolic rate (Albers, 2012; Chen & Hong, 2018; Freire-Regatillo et al., 2017; O’Connell & Hofmann, 2011; Tsuneoka & Funato, 2021). The mammalian mPOA is sexually dimorphic in volume and cell density (Gorski et al., 1980; Campi et al., 2013) and exhibits dense expression of steroid hormone receptors critical for sex differences in behavior (O’Connell & Hofmann, 2012; Xu et al., 2012). Prior studies have largely focused on how a limited set of neuroendocrine systems are associated with aggression and social status. Estrogen receptor alpha-expressing (*Esr1*+) neurons in the mouse mPOA have been implicated in driving sex-specific behaviors (Wei et al., 2018) and estrogen receptor gene expression levels in the mPOA vary with social status in females but not males (W. Lee et al., 2022; Williamson et al., 2019). Similarly, *Esr1*+ neurons in the mPOA projecting to the ventrolateral part of the ventromedial hypothalamus (VMHvl) suppress female infanticide and male aggression, whereas projections to the principal nucleus of the bed nucleus of the stria terminalis (BNSTpr) promote aggression in both sexes (Mei, Yan, et al., 2023; D. Wei et al., 2023). Given that mPOA neurons co-express diverse neuropeptides and neurotransmitters (Tsuneoka et al., 2017) and that neuropeptide and receptor co-expression is common in hypothalamic neurons (Chen et al., 2017; Romanov et al., 2017), exploring neuropeptide co-expression at the cellular level is essential to understand their role in social dominance (Miller-Crews & Hofmann, 2025).

Single-cell and single-nucleus RNA sequencing (snRNA-seq) offers unprecedented resolution to characterize cellular transcriptomes and molecular responses across conditions (Wang & Navin, 2015). These methods have demonstrated that the hypothalamus is composed of highly heterogeneous cell populations with distinct gene expression and regulatory profiles (R. Chen et al., 2017; D. W. Kim et al., 2020; Langlieb et al., 2023; S. D. Lee et al., 2019; Romanov et al., 2017, 2020). Within the mPOA, single-cell approaches have further annotated spatial molecular mapping of neuron subtypes and resolved molecularly defined cell types linked to homeostatic functions (Chung et al., 2017; Guo et al., 2024). Importantly, snRNA-seq can clarify whether sex differences arise from changes in cell number, composition, or both (Gegenhuber & Tollkuhn, 2020), and recent work has begun characterizing sex differences in neuroendocrine receptor expression in mPOA cells (Gegenhuber et al., 2022; Kaplan et al., 2025). However, across the entire mouse brain, only 21 sex-biased transcriptomic clusters have been identified, concentrated in the pallidum, striatal amygdala, hypothalamus, and hindbrain, with just 5 representing small, sex-specific clusters (Yao et al., 2023), highlighting how rare and regionally constrained sex differences in cell-type composition are, and underscoring the need to examine how sex-specific mPOA cell populations regulate social status. Previously, we have shown that outbred CD-1 male and female mice housed social groups reliably form highly linear dominance hierarchies with individuals occupying unique ranks (Milewski, Lee, Young, et al., 2025; Williamson, Franks, et al., 2016; Williamson, Lee, et al., 2016; Williamson et al., 2019). We have also found that social status is associated with coordinated endocrine and neurobiological signatures including context-dependent relationships with corticosterone/testosterone in males (Milewski et al., 2025; Williamson et al., 2019; Williamson et al., 2017) and hypothalamic gene expression profiles in both sexes (W. Lee et al., 2022; Williamson et al., 2019).

Here, we use snRNA-seq to test how sex and social status are encoded in mPOA neuronal transcriptomes. First, given that the mPOA contains distinct hormone-sensitive and sexually dimorphic neuronal subtypes, we hypothesized that sex and social status would be reflected in both the representation of specific neuronal subpopulations and the gene expression signatures within those populations, rather than appearing solely as a uniform shift across all neurons. We tested this by quantifying rank and sex effects on cluster and HypoMap-defined subtype proportions (Steuernagel et al., 2022) and using differential expression and rank-rank overlap analyses to distinguish shared versus sex-specific status-associated transcriptional profiles. Second, since hypothalamic neurons commonly express neuropeptides and peptidergic systems in the mPOA are tightly linked to social behavior and internal-state regulation, we hypothesized that social status and sex would reorganize peptidergic regulation in the mPOA through changes in peptide expression and co-expression. This was tested by comparing neuropeptide prevalence and co-expression network structure by sex and status. Overall, these analyses elucidate the neuromolecular mechanisms underlying social behavior in complex social environments across life-history contexts.

## METHODS

### Animals, Housing, and Behavioral Observations

Outbred CD-1 mice (N = 120, 60 males, 60 females) aged 7- 8 weeks were obtained from Charles River Laboratory (Houston, TX, USA) and housed at The University of Texas at Austin. The animal facility was kept on a 12/12 light/dark cycle, with white light at 2300h and red lights (dark cycle) at 1100h with constant temperature (21–24 °C) and humidity (30–50%). Upon arrival, mice were pair-housed with a same-sex cage mate in standard cages with pine shaving bedding. Standard chow and water were provided *ad libitum*. Mice were marked on the neck, back, and/or flanks with a nontoxic animal marker to display unique identification during pair and group housing social behavior observations (Stoelting Co., Wood Dale, IL, USA). After 16 days in paired housing, animals were randomly assigned into same-sex groups of six mice in two standard rat cages (35.5 x 20.5 x 14cm) covered in pine shaving bedding and connected by Plexiglass tubes with multiple enrichment objects (**Figures 1A-B**; 6 total group, 3 for each sex). Live behavioral observations began on GD1 and were conducted daily until GD20 during the dark cycle by a trained observer, as described previously (So et al., 2015; Williamson, Lee, et al., 2016). Briefly, all occurrences of agonistic (fighting/biting, chasing, mounting) and subordinate behaviors (freezing, fleeing, subordinate posture) between two individuals were recorded using either an Android or Apple device and directly uploaded to a timestamped Google Drive via a survey link. On average, 75 minutes per day of live observation was conducted per group, resulting in an average of 173 agonistic/subordinate behavioral interactions per group. All procedures were conducted with the approval of the Institutional Animal Care and Use Committee of the University of Texas at Austin (Protocol No. AUP-2019-00155).

**Figure 1.**
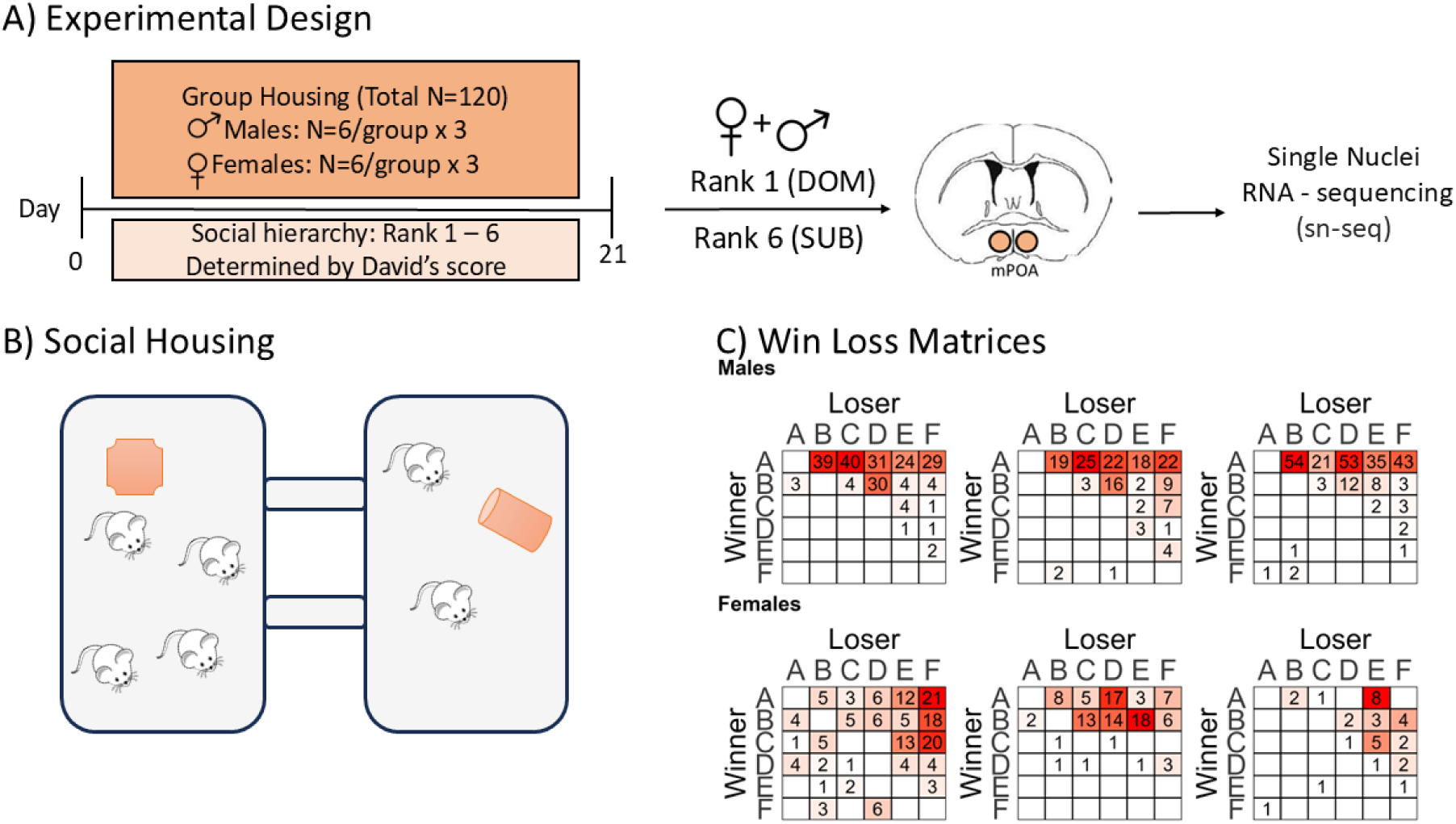
Experimental design and social dominance hierarchies. **A)** Mice were house in same-sex groups of 6 for 21 days and assigned relative ranks within their home-cage hierarchy via David’s score. The top-ranked individual (A – rank 1) was denoted ‘dominant’ (dom) and the bottom-ranked individual (F – rank 6) was denoted ‘subordinate’ (sub). The medial preoptic area of the hypothalamus was collected from three individuals of each rank and sex (12 individuals total) and processed for single-nucleus RNA-sequencing (snRNA-seq). **B)** A large, enriched cage system was used for social group housing of mice. **C)** The win-loss matrices of each social group were constructed from the results of regular home-cage behavioral observations. Color intensity is scaled to the minimum (white) and maximum (red) number of dyadic agonistic encounters recorded in each community.

### Social Behavior

Win/loss sociomatrices for each social group were created by aggregating wins and losses observed from home-cage social behavior observations across the 21 days of group housing (**Figure 1C**). Using these sociomatrices, we tested for the significance of directional consistency as previously described (Williamson et al., 2019; Williamson, Lee, et al., 2016) to assess the stability of each social hierarchy. Individual social ranks were determined by David’s Scores (DS). DS is a measure of individual dominance related to the proportion of wins to losses adjusted for opponents’ relative dominance (Gammell et al., 2003). DS was calculated after group housing was completed using the R package ‘compete’ (Curley, 2016).

### Tissue collection and processing

Midway through the light cycle on day 21 of group housing (GD21), mice were retrieved from the group housing system. Mice were weighed and euthanized via rapid decapitation. Brain tissue was dissected and immediately flash frozen in a hexane cooling bath chilled on dry ice, then stored at −80 °C until further processing. Collected whole brains were sectioned on a cryostat (Leica Biosystems, Deer Park, IL) at 200µm and the mPOA was dissected separately using a Stoelting 1 mm tissue punch (Stoelting, Wood Dale, IL). The mPOA was dissected from five coronal brain sections identified using the Allen Mouse Brain Atlas with the anterior commissure serving as the anatomical landmark, as the mPOA is located ventral to this structure (approximately +0.15mm to +0.30mm from Bregma), for a total of ten punches per individual (five per hemisphere). Female samples selected for snRNA-seq were taken from groups in which the dominant and subordinate females were both in diestrus at the time of euthanasia. All animals selected were from groups representative of typical hierarchies for their sex. Punches were pooled from three mice in each of the four experimental conditions: dominant male, dominant female, subordinate male, and subordinate female (1 pool per group, n=3/pool). The tissue punches from pooled samples weighed 16.5 mg on average.

### Nuclei isolation

We adapted a density gradient protocol for nuclei isolation (Miller-Crews & Hofmann, 2025). Briefly, all tissue samples were frozen or on ice throughout the protocol to prevent any additional transcriptional activity. Pooled tissue samples were homogenized in 1 mL of chilled Nuclei EZ Lysis Buffer (Sigma-Aldrich, USA) with 0.2 U/µl Protector RNase Inhibitor (Roche, USA) before homogenization with a chilled 7 ml Dounce homogenizer with pestle "b”. Between pooled samples, the Dounce homogenizer was cleaned with chilled 100% EtOH and chilled 1x PBS. Immediately after homogenization, nuclei were filtered through a 35μm cell strainer cap (Falcon Round-Bottom Polystyrene Test Tubes with Cell Strainer Snap Cap, 5mL Corning, USA) into a 1.5 mL tube. Nuclei were then centrifuged at 900rcf for 5 min at 4°C. The supernatant was discarded and the nuclei pellet resuspended for 1 minute in 750μl of 25% Optiprep (Sigma-Aldrich, USA), 2% BSA, and 0.2 U/µl Protector RNase Inhibitor (Roche, USA) in 1x PBS. Nuclei in this resuspension buffer were then overlaid onto 750μl of a 29% Optiprep nuclei resuspension buffer and then centrifuged at 13,000rcf for 30 min at 4°C. The supernatant was again discarded while the 1.5 mL tube was held at an angle similar to the centrifuge angle to minimize debris. The nuclei pellet was resuspended with 50μl of nuclei resuspension buffer by gently pipetting up and down. Resuspended nuclei were quickly transferred to a new 0.5ml tube with an added 50μl of nuclei resuspension buffer. One additional filtration was performed with a 35μm cell strainer cap (Corning, USA). Finally, nuclei concentration was quantified by removing a 5μl aliquot and adding to 5μl resuspension buffer with 10μg/mL DAPI. All four sample pools were processed at the same time, and this protocol took a total of ∼80 minutes.

Nuclei were counted twice per sample pool on the Countess 3 Automated Cell Counter (Invitrogen, USA) with a goal concentration between 700-1200 nuclei/μl. Additionally, samples were manually counted under a microscope using a hemocytometer and we visually confirmed nuclei quality. Samples were sent the UT Austin’s Genomic Sequencing & Analysis Facility for quality control and library construction on the 10X Genomics Chromium Next GEM Single Cell 3’ reagent kit v3.1 as per manufacturer’s instructions. The libraries were then sequenced on the Illumina NovaSeq S1 platform.

### Processing of snRNA-seq data

In total, 1,408 million reads were captured across all samples (dominant males: 390 million reads, subordinate males: 378 million reads, dominant females: 256 million reads, subordinate females: 384 million reads). Fastq files were assessed for quality control assessment with MultiQC (Ewels et al., 2016) to confirm that all reads had Phred scores above 35. The Cell Ranger 6.1.2 pipeline (Zheng et al., 2017) and the 10x Genomics prebuilt mouse reference (refdata-gex-mm10-2020-A) were used for read mapping and annotation, resulting in an average of 90.2% reads mapped to the mouse genome. Genotype demultiplexing and doublet/multiplet detection was performed using the Souporcell package in Python (Heaton et al., 2020) to identify three putative individuals per sample pools and filter for single nuclei transcriptomes. Next, we implemented the Seurat v4.4 integrated analysis pipeline (Satija et al., 2015) with reciprocal PCA and integrated the four transformed datasets with functions SCTransform v2 and IntegrateData (Choudhary & Satija, 2022). The data for each pooled sample was again filtered by selecting genes expressed in a minimum of 3 cells and selecting nuclei within a range of detected genes (nFeature_RNA) as determined by the sample pool distribution to reach the approximate targeted 10,000 nuclei per sample pool. This threshold was determined from the distribution of nuclei to reduce both background noise and doublets/multiplets. The clustering resolution of integrated expression data was determined by using Clustree (Zappia & Oshlack, 2018) to select the lowest resolution yielding a stable number of clusters (res = 0.8).

Cell type annotation was performed using the HypoMap mouse hypothalamus database (Steuernagel et al., 2022), the ScType package (Ianevski et al., 2022), and single-cell variation inference (Lopez et al., 2018). Cell type was assigned if the prediction probability was greater than 0.75 for a single broad cell type class or ‘unknown’ if less than or equal to this threshold. Neuronal nuclei were primarily classified as either inhibitory or excitatory and then assigned to the secondary hypothalamic neuronal subtypes identified in the HypoMap database (levels C7 and C66). All neuronal nuclei identified above were also re-clustered using Clustree (Zappia & Oshlack, 2018) and Uniform Manifold Approximation and Projection (UMAP) in Seurat (Satija et al., 2015) to categorize distinct expression profiles (clusters) based on differential gene expression between clusters (regardless of HypoMap subtype).

Marker genes for each cluster were identified with the FindAllMarkers function from Seurat, which classifies them based on differential expression of each gene in a given cluster compared to all other clusters. The specificity score for each cluster marker gene was calculated as the average *log_2_FC* compared to the other clusters multiplied by the relative cluster size (proportion of all neurons in that cluster) and the percent of cells in the cluster expressing the marker gene (pct1) divided by the percent of neurons in all other clusters expressing the same gene (pct2), or: specificity = avg.*log_2_FC\**(pct1/pct2)*relative cluster size. This weighted specificity score ranks marker genes for each cluster by how specific each gene is to neurons in that cluster relative to all other neurons and is offset by the proportion of neurons in the cluster to prevent nonlinear inflation of specificity scores as cluster size decreases. Importantly, weighting by cluster size does not alter the rank order of marker genes within each cluster. The top 5% of genes with the highest weighted specificity scores within each cluster were selected as cluster marker genes, yielding 413 total unique marker genes across all clusters (**Table S5**), with some genes appearing as marker genes for more than one cluster. Subsequent analyses testing for differential expression of these marker genes by social status and sex (**Figure 2D**) as well as group differences in cluster assignment are described below.

**Figure 2.**
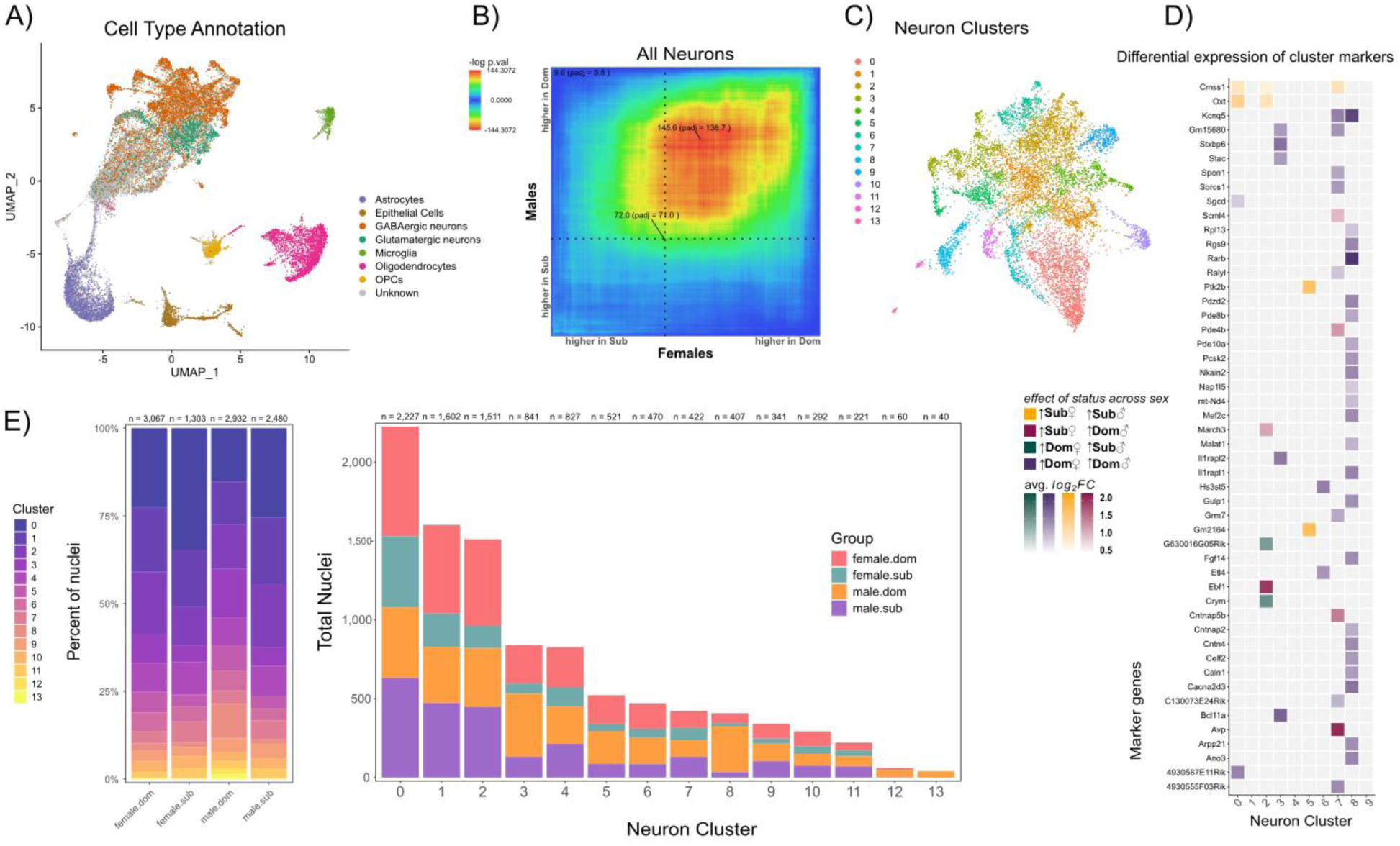
Clustering, annotation, and patterns of differential expression in neuronal nuclei. **A)** Cell types were assigned after integration and clustering of all sample pools. ScType and HypoMap cell type annotation results were combined to assign primary cell classes to all nuclei with a high level of confidence, and assignments < 0.25 confidence were labeled ‘unknown’. **B)** After filtering to neurons only, rank-rank hypergeometric overlap (RRHO) analysis was used to identify significantly enriched quadrants of genes by sex and social status. The heatmap of - log10 p-values indicates degree of enrichment, where greater expression of genes relative to the same-sex counterpart of opposite social status is represented by four quadrants: dominant males and subordinate females (<u>top left</u>), dominant males and dominant females (<u>top right</u>), subordinate males and dominant females (<u>lower right</u>), or subordinate males and subordinate females. **C)** Nonlinear dimensionality reduction via UMAP identified fourteen transcriptionally distinct clusters of neuronal nuclei. **D)** Cluster marker genes (95^th^ percentile or higher specificity scores per cluster) that were also differentially expressed by social status in both sexes (*eFDR* < 0.05 and |*log_2_FC*| > 0.2) are represented across the 10 neuron clusters that include nuclei from all individuals. Color and intensity indicate direction and magnitude of differential expression in each cluster. **E)** Proportions (<u>left</u>) and counts (<u>right</u>) of neuronal nuclei in each cluster are divided by sample pool.

### Analysis of neuron clusters and subtypes

The proportions of neurons in each cluster were compared across social status and sex with binomial GLMs (Fuess & Bolnick, 2023) and adjusted for multiple comparisons within clusters using a multivariate *t*-distribution (Hothorn et al., 2008), then adjusted with Benjamini-Hochberg FDR correction to account for comparisons across all clusters tested. For models with a significant interaction effect, the marginal effects at representative values (MER) of each interaction term component and their average marginal effects (AME) were calculated separately (Lenth et al., 2025; McCabe et al., 2022; Searle et al., 1980). Main effects are reported for models without a significant interaction term.

### Differential gene expression analyses

Differentially expressed genes (DEGs) in neurons within each cluster or primary subtype (inhibitory vs. excitatory) were identified using the limma package in R for Bayesian modeling (Law et al., 2014; Soneson & Robinson, 2018). Additionally, genotype data distinguishing the putative individuals in pooled samples was incorporated to increase statistical power. Pseudo-bulk datasets were generated by averaging expression values across neurons for each putative individual and separated by cluster and primary neuron type. Model selection was performed using the selectModel function (limma) and iteratively comparing models of differential gene expression including various combinations of interaction terms and main effects from the groupings above. Individual weights corresponding to the proportion of nuclei in each grouping contributed by an individual in a sample pool were included in the contrast matrix to account for the variable number of cells contributed by that individual to the given sample pool. The empirical false discovery rate (*eFDR*) for each gene was calculated using 10,000 permutations of sample/group assignments within the pseudo-bulk dataset by summing the number of permutations that generated a larger test statistic value (or smaller p-value) than the true data and dividing by the number of permutations: *eFDR* = Σ(permutation p-value < true p-value)/total permutations.

### Threshold-free analysis of concordance and discordance

Rank-rank hypergeometric overlap (RRHO) (Cahill et al., 2018) is a common analytic method for assessing directional patterns of DEG overlap across comparisons. We employed a recent update to this method from the RedRibbon package (Piron et al., 2023) to identify the concordant and discordant effects of social status on gene expression patterns across all neurons of each sex. Gene ontology (GO) enrichment analysis was done using ClusterProfiler v4.0 (Wu et al., 2021). Additionally, we used RRHO to analyze gene expression patterns in certain neuronal clusters or subtypes.

### Neuropeptide co-expression network analysis

A list of 127 known neuropeptide genes encoding pre-prohormone mRNAs was compiled from multiple databases (Burbach, 2010; Y. Kim et al., 2011; Seal et al., 2023) and via Ensembl (Martin et al., 2023). Next, we generated a presence/absence matrix for these neuropeptide genes across neuronal nuclei from each putative genotype across social status and sex with a threshold of two raw (not normalized) reads, or >1 raw transcript. The proportion of neuron nuclei expressing individual neuropeptides genes was compared across social status and sex with binomial GLMs as described above. Additionally, we compared the proportions of neuronal nuclei expressing multiple neuropeptides across sex and social status. The cumulative distributions of the proportions of neurons expressing 0-9 neuropeptide genes – no nuclei contained >1 raw transcript of more than 8 neuropeptide genes – were compared with a Monte Carlo two-sample Kolmogorov-Smirnov test.

Similarity between the covariance matrices generated from the normalized expression values of all 52 substantially expressed neuropeptide genes was tested between all four sample pools using methods for high dimensional data to generate HD (Chang et al., 2017), CLX (Cai et al., 2013), and LC (Li & Chen, 2012) statistics. The relative similarities between co-expression networks of the top 16 neuropeptide genes (those expressed in >1% of neurons) were broadly compared between sample pools using the Euclidean distances between eigenvalue spectra generated from their covariance matrices. We then calculated the Spearman correlations (ρ) calculated for each pair of co-expressed neuropeptide genes in the sample pool to measure specific pairwise relationships between genes. The differences between Spearman correlation values (Δρ) when comparing sample pools to each other were reported for each neuropeptide pair.

## RESULTS

### Male and female CD-1 mice placed in same-sex social group housing form stable linear dominance hierarchies

Twenty groups (n=10 males; n=10 females), each consisting of six mice, were systematically observed over 21 days to record agonistic interactions among cage-mates (**Figures 1A-B, S1**). Three groups per sex were selected for snRNA-seq, with the most dominant (dom) and most subordinate (sub) individuals from each cage included in the corresponding sex-and rank-specific sample pools for a total of four sample pools (male dom, male sub, female dom, female sub). Dyadic win–loss interactions of these three groups are visualized as sociomatrices in **Figure 1C**, showing that both males and females consistently established and maintained social dominance hierarchies within the selected groups. All groups formed stable linear hierarchies with highly significant directional consistency (all p < 0.001). Median directional consistency (DC) was 0.97 [IQR: 0.965–0.97] in males and 0.90 [IQR: 0.78–0.92] in females.

### snRNA-seq reveals expected cell type inventories

We successfully sequenced the transcriptomes of single nuclei from all four sample pools comprised of hypothalamic preoptic area tissue from 12 individuals total (n=3/social status/sex). After filtering and quality control, we identified on average 6,705 ± 178 nuclei per sample pool (female dom – 6,869; female sub – 6,848; male dom – 6,559; male sub – 6,543; **Figures S2A-D** respectively). Samples were de-multiplexed in Python using the Souporcell package to identify three putative genotypes per sample pool (Heaton et al., 2020). On average, 2,235 ± 298 nuclei were attributed to each distinct genotype. Integration and clustering of all four sample pools yielded 22 common cell clusters across all samples and genotypes (**Figure S3**). Cell type annotation was based primarily on the HypoMap reference transcriptomes (Steuernagel et al., 2022) and then compared to ScType cell type assignments (Ianevski et al., 2022). We recovered consistent proportions of seven major hypothalamic cell types: inhibitory (GABA) neurons, excitatory (GLU) neurons, oligodendrocyte progenitor cells (OPCs), oligodendrocytes, astrocytes, microglia, and epithelial cells (**Figures 2A and S4A-D**). These were confirmed by cross-referencing their corresponding cluster marker genes. All neurons were identified as either excitatory or inhibitory and were isolated from other cell types based on this assignment. The proceeding analyses and their results are limited to the population of neuronal nuclei only.

### Gene expression differences between dominant and subordinate individuals are remarkably concordant across sexes

To assess broad patterns of expression across all neuronal cells, ordered gene lists were first constructed by ranking genes via magnitude and direction (signed *log_2_FC*) of differential expression across social status and within each sex. Rank-rank Hypergeometric Overlap (RRHO) analysis was performed to identify concordant and discordant effects of social status on gene expression patterns in a threshold-free manner. The intersection of the two sex-specific gene lists was most significantly enriched by genes with higher expression in all dominant individuals (up-up) compared to same-sex subordinates (up-up – 852 genes higher in all dominants, *p.adj* < 0.001) (**Figure 2B**). The opposite concordant quadrant was also significantly enriched (down-down – 953 genes higher in all subordinates, *p.adj* < 0.001). The concordant effects of social rank indicate the presence of common status-related expression shifts in dominant and subordinate individuals of both sexes. The results of Gene Ontology (GO) enrichment analysis for biological processes (BP) on these concordant gene sets are summarized in **Figure S5A-B**. Briefly, dominant animals exhibited higher expression of genes associated with dendrite development, synapse assembly, regulation of membrane potential, and learning/cognition/memory. In comparison, subordinate individuals had relatively higher expression of genes associated with regulation of post-translational protein modification and protein stability, translation at synapse, and regulation of intracellular protein transport. Neither discordant gene set was enriched by overlapping genes (higher in dominants or subordinates of one sex and lower in the other, up-down = 2 genes, *p.adj* = 1; down-up = 3 genes, *p.adj* = 0.680) (**Table S1**), meaning that there were 361 times more concordant genes (1805) than discordant genes (5) (Binomial Test: 95% CI = [0.994, 1.000], p < 0.001).

We also used RRHO to analyze excitatory and inhibitory neurons separately to determine whether this pattern of concordance depends on primary neuron subtype. We found that genes in the up-up quadrant were similarly enriched in both GABAergic (up-up = 753 genes, *p.adj* < 0.001) and glutamatergic (up-up = 1428 genes, *p.adj* < 0.001) neurons (**Figure S5C**). The opposite concordant quadrant (down-down) was also significantly enriched by overlapping genes in both neuron types (GABA = 969 genes, *p.adj* < 0.001; GLU = 995 genes, *p.adj* < 0.001). Additionally, the up-down quadrant (higher in dominant males and subordinate females) of overlapping genes in glutamatergic neuronal nuclei was also significantly enriched (up-down = 4 genes, *p.adj* < 0.001) and the opposite discordant quadrant (down-up – higher in subordinate males and dominant females) approaches significance in GABAergic neurons (down-up = 8 genes, *p.adj* = 0.071) (**Table S1**).

### Relative proportions of neuronal sub-classifications vary by sex and status

Due to the documented heterogeneity of neuronal subtypes in the mPOA (Steuernagel et al., 2022), we used independent graph-based clustering to identify fourteen neuron clusters with statistically distinct gene expression profiles (**Figures 2C and S6A-B**). Clusters #12 and #13 were composed of nuclei from dominant individuals alone, and only clusters #0-9 contained nuclei from all 12 putative genotypes (**Figures 2D and S6C-D**). Although each cluster contained varying proportions of excitatory and inhibitory neurons, clusters #0-9 were comprised of both primary neuron sub-classes (**Figures S6E and S7A**). On average, ∼26% of neurons were classified as excitatory and ∼74% were classified as inhibitory, and analysis with FDR-corrected binomial GLMs revealed no significant effects of sex, social status, or the interaction of sex and social status on excitatory/inhibitory neuron proportions (*FDR* > 0.05 for all comparisons) (**Table S2**). Excluding clusters #10-13, there was a significant interaction of sex and social status on the proportion of neurons in clusters #1, #2, #3, and #8 (1: β = 0.667 ± 0.117, FDR < 0.001; 2: β = 0.994 ± 0.126, FDR < 0.001; 3: β = -0.525 ± 0.178, FDR < 0.001; 8: β = -1.671 ± 0.326, FDR = 0.010). Dominant males had greater proportions of cluster #3 and cluster #8 neurons compared to subordinate males, while subordinate males had relatively greater proportions of cluster #1 and cluster #2 neurons (**Table S3**). Compared to subordinate females, dominant females had greater proportions of cluster #2 and cluster #3 neurons, and there was no difference between social ranks for female neurons in clusters #1 and #8 (**Figure 2E**). For both males and females, dominant mice had fewer cluster #0 and cluster #7 neurons relative to subordinates (0: β = 0.616 ± 0.050, FDR < 0.001, 7: β = 0.458 ± 0.101, FDR < 0.001), although dominant mice had greater relative proportions of clusters #5 and #6 neurons (5: β = -0.688 ± 0.104, FDR < 0.001, 6: -0.415 ± 0.104, FDR < 0.001). In dominant individuals, but not subordinates, males also had significantly more neurons than females in cluster #3. There was a main effect of sex only for clusters #0 and #9: females had significantly greater proportions of neuronal nuclei from cluster #0 neurons than males (0: β = -0.465 ± 0.050, FDR < 0.001), while males overall had higher proportions of neurons in cluster #9 (9: β = 0.332 ± 0.116, FDR = 0.021).

We determined that cluster assignments did not solely reflect differences in predicted spatial origin (i.e. hypothalamic subregions), although the predictions of regional origin based on HypoMap sample data were associated with a high degree of uncertainty (**Figure S7D**). Additionally, we noted that that cluster assignment was not based on the neuronal subtype assignments established in HypoMap (Moffitt et al., 2018a; Steuernagel et al., 2022). Each cluster contained varying proportions of multiple neuronal subtypes classified in HypoMap, and none of these were exclusive to any particular cluster (**Figure S8A-C**). Again, FDR-corrected binomial GLMs were used to analyze differences in the proportions of these neuronal subtypes (**Table S4**). Of the 21 most common HypoMap subtypes (containing > 1% of neurons in each sample pool), there was a main effect of social status wherein dominant males and females both had greater proportions of *Samd3* (sterile alpha motif containing domain 3) glutamatergic neurons while subordinates had more *Lhx1* (LIM homeobox 1) GABAergic neurons (*Samd3*: β = -0.629 ± 0.117, FDR < 0.001, *Lhx1*: β = 0.490 ± 0.182, FDR = 0.019). The main effects of sex and social status were independently significant for both *Sst* (somatostatin) and *Pmch* (pro-melanin concentrating hormone) glutamatergic neurons, of which subordinates overall had greater proportions compared to dominants (*Sst*: β = 1.094 ± 0.181, FDR < 0.001; *Pmch*: β = 1.145 ± 0.241, FDR < 0.001) and females had overall greater proportions than males (*Sst*: β = - 1.259 ± 0.0.193, FDR < 0.001; *Pmch*: β = -1.914 ± 0.290, FDR < 0.001). In addition, there were significant main effects of sex and social status on the proportions of *Vipr2* (vasoactive intestinal peptide receptor 2) GABAergic neurons, which were higher in all females regardless of social status (*Vipr2*: β = -0.398 ± 0.116, FDR = 0.003) and in all subordinates regardless of sex (*Vipr2*: β = 0.285 ± 0.116, FDR = 0.041) (**Figure S8B**).

The remaining group differences in the neuronal subtype assignments were predominantly attributed to the interaction of sex and social status, indicating that the effect of status often varied depending on sex and vice versa. All dominant individuals had higher proportions of *Lhx8* (LIM homeobox 8), *Meis2* (Meis homeobox 2), and *Chat* (choline O-acetyltransferase) GABAergic neurons (*Lhx8*: β = -0.406 ± 0.159, FDR = 0.034; *Meis2*: β = - 2.412 ± 0.309, FDR < 0.001; *Chat*: β = -1.347 ± 0.419, FDR = 0.007) and dominant males had markedly more neurons of these subtypes than any other group. Dominant females in particular had the highest proportions of *Lef1* (lymphoid enhancer binding factor 1) and “mixed” GABAergic neurons (*Lef1*: β = 0.714 ± 0.223, FDR = 0.008; mixed GABA: β = 0.518 ± 0.109, FDR < 0.001) as well as *Gpr149* (G protein-coupled receptor 149) glutamatergic neurons (β = 1.491 ± 0.424, FDR = 0.003), although subordinate females had the greatest proportion of *Onecut2* (One cut homeobox 2) GABAergic neurons (β = 0.555 ± 0.190, FDR = 0.012). In contrast, subordinate males had the highest proportion of *Nts* (neurotensin) GABAergic (β = 0.612 ± 0.142, FDR < 0.001) and *Trh* (thyrotropin-releasing hormone) glutamatergic neurons (β = 1.203 ± 0.204, FDR < 0.001). While subordinates of both sexes had increased proportions of *Caprin2* (caprin family member 2) glutamatergic neurons, the difference between ranks was greatest in males (β = 1.029 ± 0.348, FDR = 0.014, **Table S4 and Figure 8B**).

### Status-dependent differential gene expression of cluster marker genes is variable across neuron clusters

As the distribution of neuronal nuclei across independent clusters primarily reflected differences in sex and social status, we next sought to identify the marker genes for each cluster to generate a candidate gene set that we then tested for differential expression by social condition across all neuronal nuclei. Marker genes for each cluster were distinguished by significantly higher expression in that cluster compared to all others, and the top 5% of genes with the greatest weighted specificity scores were selected as the marker genes for each cluster (**Figure S7B and Table S5**). An inverse relationship exists between cluster marker specificity and cluster size, which was dampened but not completely eliminated with the use of weighted specificity measures. 50 of the 300 marker genes identified for clusters #0-9 were differentially expressed by social status in male and female neurons in at least one of these clusters, 38 of which were differentially expressed (DE) in at least one cluster they were not a marker for, revealing that expression of these genes is variable across all neurons and is significantly associated with social status(**Figure 2D**). Cluster #8 contained the greatest number of marker genes (of these 50) that were DE by social status in both sexes. Within this cluster, all 23 genes exhibited higher expression in dominant individuals compared to subordinates. Of these, the voltage-gated potassium channel (subfamily Q member 5) gene *Kcnq5*, the retinoic acid receptor beta gene *Rarb*, and *Avp* (arginine vasopressin) were the top 3 marker genes with the highest average *log_2_FC* values between social ranks (*Rarb* avg.*log_2_FC* = 2.167; *Kcnq5* avg.*log_2_FC* = 1.989; *Avp* avg.*log_2_FC* = 2.082).

### Differential gene expression across clusters is more directionally consistent in females

To investigate the expression patterns of genes with the most widespread status-dependent differential expression in neurons, we identified genes that were DE by social status in at least half of the 10 largest clusters (#0-9) and found that 158 genes were DE in females but only 6 genes were DE in males. Within this gene set, only ∼30% of differential expression in males was associated with dominant status (**Table S6**). Statistics for all DE genes (DEGs) can be found in **Table S7**. Notably, the top three genes DE by social status for males in the most clusters were *Trh*, *Oxt* (oxytocin), and *Avp*, which were consistently higher in subordinates, while the most broadly DE genes across clusters in females were *Cwc22* (CWC22 spliceosome associated protein), *Pmch*, and *Ralyl* (RALY RNA binding protein like) and were all higher in dominant animals (**Figure S7C**). Overall, females had over seven times as many DEGs as their male counterparts, which were expressed with greater directional consistency across clusters.

### Oxytocin transcript abundance is the primary driver of variation in neuropeptide gene co-expression patterns across sex and social status

To identify differentially expressed genes while controlling for differences in neuron composition, ensuring that DEGs reflect higher expression per neuron rather than merely a higher number of neurons expressing them, we next investigated the differential expression of genes across all neurons by weighting model terms based on neuron cluster and primary subtype (GABAergic/glutamatergic). For both males and females, *Oxt* was among the top 10 DEGs in all neurons (ranked by *log_2_FC* and *eFDR*) with higher expression in subordinates compared to their dominant counterparts (male dom – male sub: *log_2_FC* = -1.486, *eFDR* < 0.001; female dom – female sub: *log_2_FC* = -1.003, *eFDR* < 0.001) (**Figure S9A-B**). The number of distinct and shared genes that are differentially expressed by social status is depicted in **Figure S9C**. In males, *Avp* was among the top 10 DEGs that were overall higher throughout all subordinate neurons (male dom – male sub: *log_2_FC* = -2.215, *eFDR* < 0.001). This effect of social status on *Avp* expression was shared by females only in excitatory neurons and not inhibitory neurons (GLU female dom – GLU female sub: *log_2_FC* = -0.681, *eFDR* = 0.0256; GABA female dom – GABA female sub: *log_2_FC* = -0.140, *eFDR* = 0.5647). In female cluster #7 neurons specifically, *Avp* was differentially expressed in the opposite direction (cluster #7 female dom – female sub: *log_2_FC* = 1.056, *eFDR* = 0.0197) (**Figure 3A**). Otherwise, the expression patterns of *Oxt* and *Avp* transcripts were highly concordant across neuron clusters and subtypes (**Figures S11 and S12**).

**Figure 3.**
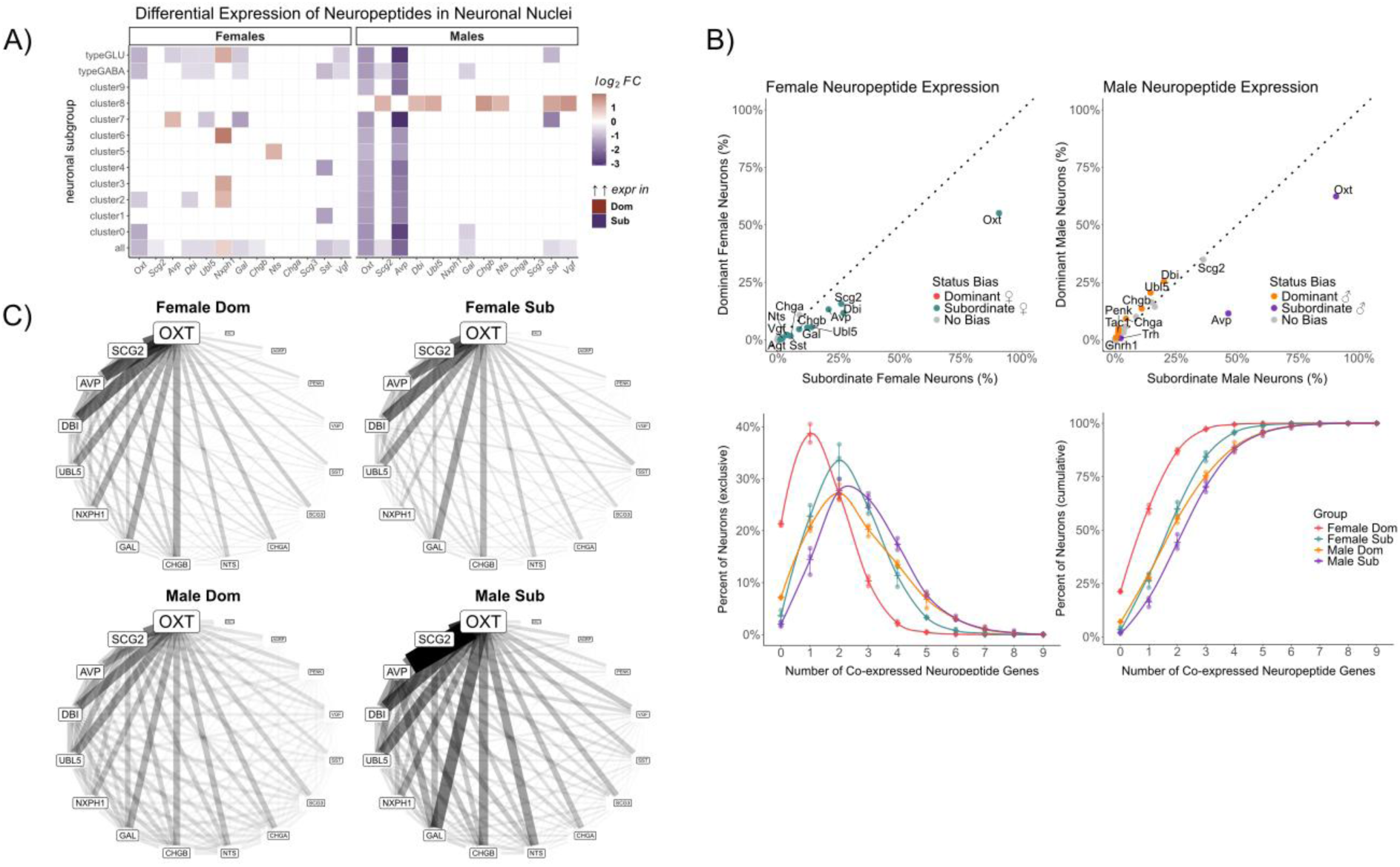
Differential expression and co-expression patterns of neuropeptide (NP) genes in neuronal nuclei. **A)** Magnitude and direction of differential expression of NP genes by social status, across neuronal subtypes and sex (eFDR < 0.05). NP genes are ordered left to right by relative abundance (percent of neurons expressing >1 read). **B)** NP gene co-expression network using a presence-absence matrix based on neuronal nuclei expressing >1 read in at least 1% of nuclei and scaled for quantity of nuclei per sample. Line width and shade intensity represents the strength of overlap across all samples; gene label size indicates the proportion of neuronal nuclei expressing >1 read, with genes arranged counterclockwise in descending order. **C)** Relative proportions of nuclei in each group expressing >1 read of all NPs for females (<u>upper left</u>) and males (<u>upper right</u>). Color indicates a statistically significant effect of social status (binomial GLM, p < 0.05) and the direction of proportion bias (greater in either dominant or subordinate animals). The raw distribution (<u>lower left</u>) and cumulative distribution (<u>lower right</u>) of NP gene co-expression (>1 read of each gene) in all neuronal nuclei are colored by sex and social status with points representing each individual in the sample pool.

To investigate the expression and co-expression levels of additional neuropeptide genes (Miller-Crews & Hofmann, 2025), we first compiled a candidate gene set from multiple databases (Burbach, 2010; Y. Kim et al., 2011; Seal et al., 2023) and via Ensembl (Martin et al., 2023) to produce a curated gene list of 127 known neuropeptide genes encoding pre-prohormone mRNAs. To test our hypotheses regarding neuropeptide co-expression, we then filtered neuronal nuclei to include only those expressing any of the 127 candidate neuropeptide genes and found that 52 genes from this neuropeptide list were expressed at levels >1 transcript in at least one neuron in our sample. The three most broadly expressed neuropeptide genes were *Oxt*, *Scg2* (secretogranin II), and *Avp*, which were expressed in 71%, 28%, and 22% of neurons respectively. Of the 9,782 total neuronal nuclei transcriptomes that were analyzed, 7,868 (80%) of them expressed >1 read of at least one of these three genes (*Oxt*, *Avp*, *Scg2*) (**Figure 3B)**. Binomial GLMs used to compare the proportion of neurons in each sample pool expressing any neuropeptide transcript yielded a strong main effect of social status on the proportion of *Oxt*+ neurons (social status: *β* = 1.873 ± 0.062, *FDR* < 0.001); 91% of neurons each from subordinate females and subordinate male neurons were *Oxt*+, which differed significantly from their dominant counterparts by 36% and 28% respectively (male dom – 55%, female dom – 63%). A similar effect was observed in *Avp*+ neurons, although to a greater extent in males than females: 46% of subordinate male neurons expressed >1 *Avp* transcript while only 12% of dominant male neurons did, and 21% of subordinate female neurons compared to only 13% of dominant female neurons (sex:status interaction: *β* = 1.369 ± 0.112). The proportions of *Scg2*+ neurons did not differ significantly between dominant and subordinate males (male dom – 35%, male sub – 36%) but were significantly higher in subordinate females compared to dominant females (female dom – 16%, female sub – 26%) (sex:status interaction: *β* = -0.559 ± 0.099, *FDR* < 0.001). All statistics for these binomial GLMs including interaction terms and the main effects of sex and social status can be found in **Table S8**.

We next explored co-expression of neuropeptide genes across neuronal nuclei. We found that on average, neurons expressed 2.21 neuropeptide genes, and the patterns of this co-expression differed across conditions. The distribution of neuropeptide co-expression varied significantly across both sex and social status; the lowest levels of co-expression were found in dominant females (average neuropeptides per neuron = 1.35), and subordinate males demonstrated the highest (average neuropeptides per neuron = 2.87) (**Figure 3B**). All four distributions were significantly different from each other (p < 0.001), although dominant males and female subordinates were the least different (**Table S9**). Covariance matrices were constructed from the normalized expression values of the top 16 neuropeptide genes (those expressed in >1% of neurons) in each sample pool. The Euclidean distances between eigenvalue spectra for each sample pool’s co-expression matrix were used to assess relative similarity between neuropeptide gene co-expression networks. Dominant males and subordinate females shared the greatest similarity, while dominant and subordinate males were the least similar (**Figure S10A**). Pairwise comparisons of these covariance matrices indicated that all neuropeptide co-expression networks were statistically different from each other (p < 0.001) (**Figure S10B**). The strength and degree of co-expression between the top 16 neuropeptides is depicted in **Figure 3C**. Pairs of genes driving these group differences were identified by calculating their Spearman correlations (ρ) and taking the difference for each pairwise comparison. Of these pairwise relationships, the largest difference in neuropeptide co-expression between dominant and subordinate animals was attributed to the stronger correlation between *Oxt* and *Avp* expression levels in subordinates. Within neuropeptide co-expression networks, increased *Oxt*-*Avp* co-expression was the primary driver of network dissimilarity between dominants and subordinates of both sexes (female dom – female sub: Δρ = -0.112, male dom – male sub: Δρ = -0.191) and the sex difference between subordinate males and females (female sub – male sub: Δρ = -0.102) such that subordinate males had increased *Oxt-Avp* co-expression compared to subordinate females. However, this difference was largely diminished between dominant males and females (female dom – male dom: Δρ = -0.022) (**Figures S10C-F**).

### Neurons expressing neuropeptide transcripts showed concordant patterns of sex- and status-specific gene expression between neuronal clusters and subclasses

In addition to the differences between social ranks in neuron proportions expressing these neuropeptide transcripts, there were large differences in the co-expression patterns of the top three neuropeptide genes (*Oxt*, *Avp*, and *Scg2*). These are shown independently, as percentages of each sub-population expressing a particular neuropeptide (**Figure 4A**), and in combination, as the proportion of all neurons expressing at least one of these neuropeptide genes (**Figure S11A**). In dominant females, 71% of all *Oxt*+ neurons expressed *Oxt* alone, which was much higher than other groups (female dom – 71%; female sub – 58%; male dom – 57%; male sub – 34%; sex:status interaction: *β* = -0.779 ± 0.089, *FDR* < 0.001). In addition to expressing the greatest proportion of *Oxt+* neurons without *Avp* or *Scg2* co-expression, dominant individuals of both sexes displayed a distinct pattern of increased *Avp-*only (sex:status interaction: *β* = 1.114 ± 0.320, *FDR* = 0.003) and *Scg2*-only neurons (N.S. sex:status interaction: *β* = -0.413 ± 0.332, *FDR* = 0.501; status: *β* = -1.989 ± 0.151, *FDR* < 0.001; sex: *β* = 0.684 ± 0.095, *FDR* < 0.001). Of all neurons expressing at least one of *Oxt/Avp/Scg2*, subordinates had the highest proportions co-expressing all three top neuropeptide genes (male sub – 16%, female sub – 5%, male dom – 3%, female dom – 2%), and males had overall greater than females (N.S. sex:status interaction: *β* = 0.442 ± 0.228, *FDR* = 0.124; status: *β* = 1.663 ± 0.101, *FDR* < 0.001; sex: *β* = 1.052 ± 0.108, *FDR* = < 0.001) (**Table S10**). Individual proportions for each putative genotype in the sample pools are depicted by cluster in **Figure S11B**.

**Figure 4.**
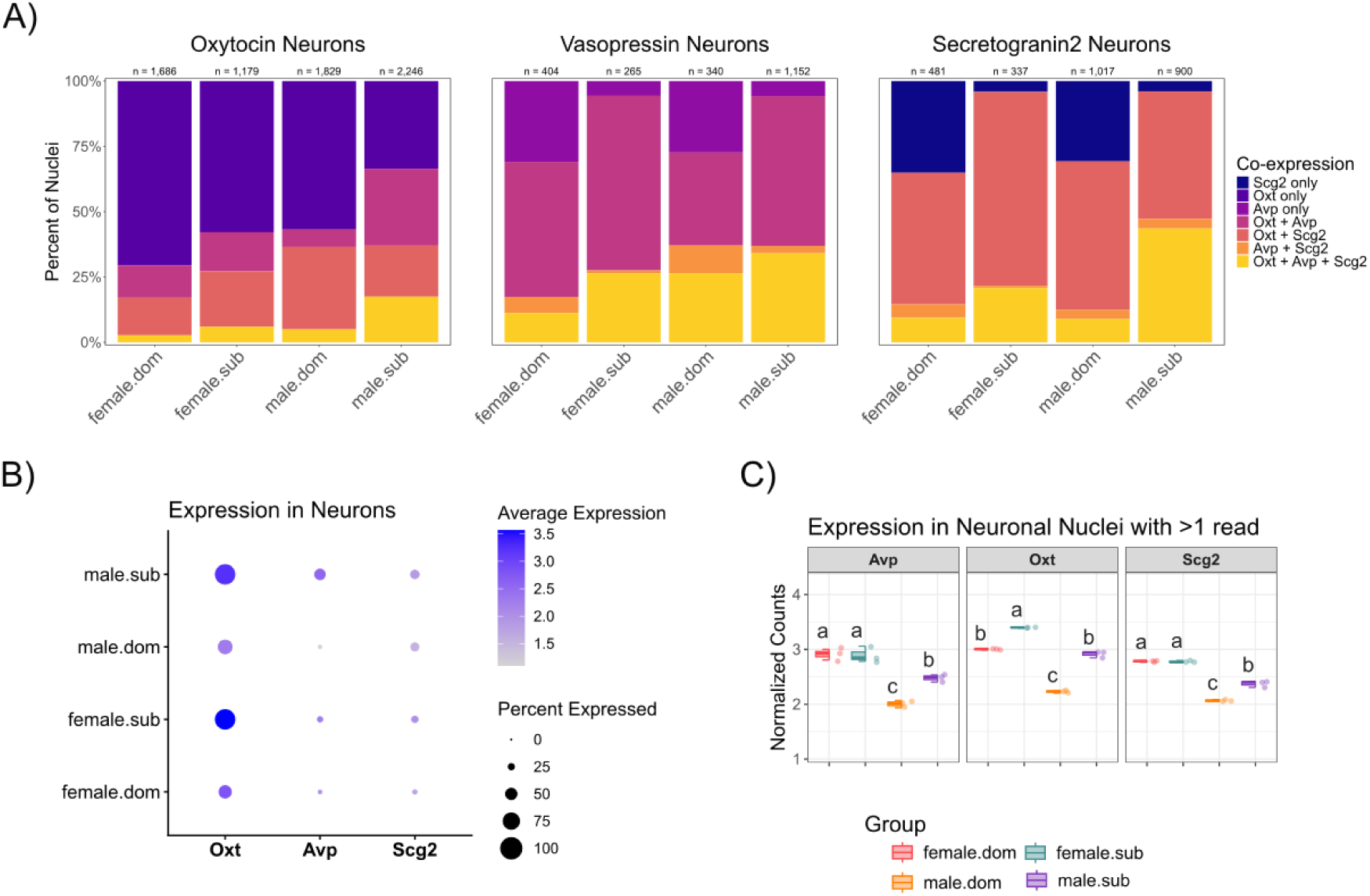
Expression and co-expression levels of the three most abundantly expressed NP genes (percent of neurons expressing >1 read). **A)** The percent of oxytocin-expressing (<u>left</u>), vasopressin-expressing (<u>center</u>), and secretogranin 2-expressing (<u>right</u>) neurons in each sample pool that express any combination of these three genes (>1 read each). **B)** The average normalized expression levels of the top three NP genes in all neuronal nuclei are indicated by color, and size indicates the percentage of all neuronal nuclei in which they are expressed (>1 read). **C)** Box plots with points representing individuals within the sample pool and their average normalized expression levels of the top three NP genes exclusively in neuronal nuclei with >1 read of that gene. Assignment to different letters indicates a significant difference between two groups.

The mean normalized expression levels in all neuronal nuclei range from 2.202 to 3.403 for *Oxt*; 1.936 to 3.060 for *Avp*; and 2.054 to 2.798 for *Scg2* (**Figure 4B**). After excluding neurons that express less than two reads (prior to normalization of expression values) of each neuropeptide gene, the effects of sex and social status on normalized expression levels remained; subordinate individuals overall had higher expression of *Oxt* in all neurons that expressed it, and females overall had increased expression of *Oxt* relative to males (all female – all male: *log_2_FC* = 0.45, *FDR* < 0.001; all dom – all sub: *log_2_FC* = -0.47, *FDR* < 0.001). Female *Avp*+ neurons also contained higher expression of *Avp* than males but with no effect of social status in females, while subordinate male *Avp*+ neurons expressed *Avp* at significantly higher levels than their dominant counterparts (all female – all male: *log_2_FC* = 0.37, *FDR* < 0.001; male dom – male sub: *log_2_FC* = -0.18, *FDR* < 0.001). A similar effect was observed in *Scg2*+ neurons: higher expression of *Scg2* occurred in female neuronal nuclei compared to male nuclei, for which there was an independent effect of social status (all female – all male: *log_2_FC* = 0.34, *FDR* <0.001; male dom – male sub: *log_2_FC* = -0.16, *FDR* <0.001) (**Figure 4C**).

These patterns were consistent across all neuron clusters with complete sample representation (clusters #0-9), except for cluster #7 regarding *Avp* expression, which was not significantly different between male dominant and subordinate mice (male dom – male sub: *log_2_FC* = -0.033, *FDR* = 1.000) (**Figure S11B-C**). When separated by HypoMap-based neuronal subtype classifications, the expression levels and neuronal proportions were again highly consistent regardless of subtype, except for *Caprin2* glutamatergic neurons (**Figures S12A-B**). This was the only HypoMap subtype in which dominant individuals exhibited similar expression levels of *Oxt* and *Avp* in neurons that expressed them and also displayed similar proportions of these neurons compared to their subordinate counterparts (%*Oxt* all dom – all sub: *β* = 0.474 ± 0.203, *FDR* = 0.103; %*Avp* all dom – all sub: *β* = 0.474 ± 0.203, *FDR* = 0.104). *Scg2* expression was relatively stable across neuron sub-classifications, with a significant main effect of social status in all subtypes except for *Caprin2* glutamatergic neurons (%*Scg2* all dom – all sub: *β* = 0.474 ± 0.203, *FDR* = 0.103) (**Figure S12C**).

### Concordant and discordant patterns of status-related gene expression in steroid receptor-expressing neurons

Given that co-expression of multiple neuropeptide and steroid receptor genes is common in mPOA neurons (Tsuneoka et al., 2017), we next investigated the gene expression patterns of steroid receptor-expressing neurons across sex and social status. Overall, we found that five steroid receptor genes (*Esr1*, *Ar*, *Pgr*, *Nr3c1*, and *Nr3c2*) were present above at levels >1 transcript in >1% of neuronal nuclei. Transcripts for the cognate receptors of *Oxt*, *Avp*, and *Scg2* were not present at appreciable levels (all <0.5% of neurons) in our sample. *Nr3c2* was expressed in the highest proportion of neurons (9% overall), and sex accounted for the most variability in *Nr3c2+* neuron proportions (N.S. sex:status: *β* = -0.206 ± 0.127, *FDR* = 0.372, status: *β* = -0.290 ± 0.050, *FDR* < 0.001, sex: *β* = 1.185 ± 0.058, *FDR* < 0.001) (**Figure S13A**). There was also a significant main effect of social status on the proportions of neuronal nuclei expressing *Esr1+*, *Ar+*, *Pgr+*, and *Nr3c1+,* but sex was consistently the strongest predictor of all five neuron proportions (**Table S11**). The results of RRHO analyses of gene expression patterns in all five subpopulations of neurons were broadly consistent with RRHO results across all neuronal nuclei, wherein sex-specific gene sets were strongly enriched with concordantly expressed genes associated with dominant status (**Figures S13B-F, Table S12**) The top GO BP terms associated with these gene sets are found in **Table S13**.

## DISCUSSION

In the present study, we applied single-nucleus RNA sequencing (snRNA-seq) to investigate how sex and social status shape neuronal gene expression patterns within the mPOA, a critical hypothalamic hub for social and reproductive behaviors (Mei, Osakada, et al., 2023). We have previously shown that both male and female CD-1 mice form highly linear social dominance hierarchies (So et al., 2015; Williamson et al., 2019). Across nearly 40,000 nuclei, we captured seven major hypothalamic cell types with a high degree of confidence, including inhibitory (GABAergic) and excitatory (glutamatergic) neurons, astrocytes, microglia, oligodendrocytes, oligodendrocyte progenitor cells, and epithelial cells, at proportions consistent with prior studies (Moffitt et al., 2018a; Steuernagel et al., 2022).

We resolved fourteen distinct subpopulations of neurons (clusters #0–13), ten of which contained nuclei from all individuals across sex and status groups. These clusters included both inhibitory and excitatory neurons, aligning with the known cellular composition of the mPOA (Hashikawa et al., 2017; D. W. Kim et al., 2025; Moffitt et al., 2018a). Cluster identities were robust to technical or regional artifacts, as HypoMap-based spatial predictions and subclassifications within each cluster remained consistent across groups. Several clusters exhibited strong sex-by-status interactions in composition, indicating that cell population variability is associated with phenotypic predictors rather than anatomical bias or batch effects. This approach revealed both shared and divergent molecular signatures of dominance and subordination in both sexes, demonstrating that hypothalamic neuronal transcriptomes are dramatically altered by social experience.

### Broad transcriptional concordance across dominant males and females

We used RRHO analysis to test whether there is a shared transcriptional signature of dominance between sexes and identified strong transcriptomic concordance between dominant males and females, a finding that is consistent with prior transcriptomic profiling of dominant and subordinate cichlid fish (Renn et al., 2008, 2016). Although subordinate males and females shared a greater number of genes with higher expression, the pattern of status-associated gene expression generated a markedly weaker signal in subordinate animals. The concordant genes associated with dominance were enriched for biological pathways associated with synaptic plasticity, learning, ion transport, and protein localization to the synapse. Consistent with findings in social fish species (Teles et al., 2016), we have previously shown that mice undergoing transitions in social status also exhibit changes in plasticity-related genes in the mPOA and medial amygdala (Milewski et al., 2025; Williamson et al., 2017). Our present results demonstrate that social dominance is associated with neuron-wide upregulation of synaptic function genes in both sexes, suggesting that across vertebrates socially dominant individuals may sustain elevated neuronal responsiveness through a conserved molecular mechanism.

We did not observe discordance in the direction of status-associated gene expression differences across sexes. This indicates that sex differences in status-related expression are driven primarily by divergent gene regulatory pathways rather than opposing regulation of the same genes. Indeed, our investigation of sex-specific differential expression between dominant and subordinate neuronal transcriptomes revealed that females exhibited a roughly sevenfold increase in the number of status-associated differentially expressed genes (DEGs) compared to males. Further, this differential expression exhibited markedly greater directional consistency in females, as nearly all DEGs were more highly expressed in dominant females. This finding is substantiated by prior work documenting widespread sex differences in socially and hormonally induced gene expression profiles (Brivio et al., 2020) in addition to evidence that female hypothalamic (and related) cell populations are more transcriptionally labile than in males (Fan et al., 2020). Also consistent with prior work suggesting sex-specific network modulation in the hypothalamus (Mozhui et al., 2012), DEGs between dominant and subordinate male neurons in our study displayed a comparatively more balanced pattern of up- and down-regulation than in females. The relatively more broad and diffuse, yet directionally consistent, pattern of status-associated differential expression in female neurons may reflect the qualitatively distinct behavioral strategies supporting the maintenance of hierarchical relationships in same-sex social groups (McCarthy et al., 2017; O’Connell & Hofmann, 2011; Renn et al., 2008).

### Cluster-specific effects of social status and sex in the mPOA

Our findings suggest that social status is a primary determinant of mPOA neuronal composition, with rank predicting shifts in cluster representation in a sex-dependent manner. In males, dominant individuals had greater proportions of cluster #3 and cluster #8 neurons, whereas subordinates had relatively more neurons in clusters #1 and #2, indicating that status is associated with a redistribution of specific neuronal populations. In females, dominant individuals similarly showed increased representation of cluster #3, consistent with a partially shared effect of status across sexes. Conversely, subordinate females had fewer neurons in cluster #2 and displayed no differences by rank in clusters #1 or #8, highlighting the sex-specific sensitivity of these neuronal populations to the social environment. These data suggest that social dominance drives coordinated changes in the relative abundance of transcriptionally distinct neuronal subgroups in the mPOA to support rank-appropriate behavioral and physiological responses.

To further investigate patterns of traditionally identified DEGs (from pseudo-bulk analysis) in neurons, we cross-referenced the set of status-associated DEGs overlapping in both sexes with the marker genes for all neuron clusters. In males and females, we saw that 38 of the top marker genes for clusters #0-9 were also differentially expressed by social status in at least one additional cluster, supporting the interpretation that cluster assignment reflects true differences in neuronal population identity driven by sex and social status rather than technical or batch effects. We found that neuron cluster #8 contained the greatest number of marker genes (23 of 50) that were differentially expressed by social status in both sexes. These genes were consistently higher in all dominant individuals, including *Rarb, Meis2,* and *Kcnq5*. The strong dominance-associated signal of elevated *Rarb* expression suggests that status-mediated transcriptional changes in mPOA neurons are regulated through retinoid signaling in dominant males and females. *Rarb* encodes retinoic acid receptor-β, a nuclear receptor that links local retinoic acid availability to changes in gene expression in the adult hypothalamus (Imoesi et al., 2019). Retinoic acid signaling has been implicated in synaptic plasticity and the stabilization of synaptic function, including homeostatic regulation at inhibitory synapses (Zhong et al., 2018). There is also increasing evidence that retinoic acid signaling regulates biological rhythms including feeding, sleep and energy balance (Ransom et al., 2014), all of which are shifted in dominant individuals (W. Lee et al., 2022; Milewski, Lee, Young, et al., 2025). In parallel, both dominant males and females exhibited significantly higher proportions of HypoMap-classified *Meis2* GABAergic neurons and overall increased neuronal *Meis2* expression compared to subordinates. *Meis2* is a transcription factor with established roles in specifying inhibitory neuron identities, consistent with these cells being positioned to integrate endocrine signals with local inhibitory network function (Dvoretskova et al., 2024). Additionally, *Kcnq5* encodes a voltage-gated potassium channel (K_v_7.5) critical for regulating neuronal excitability and promoting stabilization of neuronal firing rates that is inhibited by M1 muscarinic receptor activation (Delmas & Brown, 2005; Jiang et al., 2015; Kosenko et al., 2012). In dominant males, the proportion of *Scg2*+ nuclei in cluster #8 was also higher than in subordinate males, a difference that was absent when comparing the proportions of all neurons. It is currently unknown how *Scg2* may influence hypothalamic inhibitory circuits, but it is notable that upregulation of *Scg2* expression is required to facilitate experience-dependent plasticity of inhibitory hippocampal circuits (Yap et al., 2021). Taken together, it is plausible that these attributes of cluster #8 neurons may contribute to the stabilization of dominance-related phenotypes and behavioral state regulation by modulating inhibitory circuit plasticity. Previous work establishing that dominance is associated with increased M1 receptor activation in the prelimbic cortex and heightened expression of genes regulating cholinergic signaling in the amygdala (Chen et al., 2024; Milewski et al., 2025) also indicates a potential role of social status in regulating excitation-inhibition balance.

Another highly specific marker gene for cluster #8, *Crym* (crystallin mu), was strongly overexpressed throughout the entire population of dominant male neurons and was among the top ten DEGs in both males and females. *Crym* encodes a cytosolic thyroid hormone-binding protein that regulates the availability of triiodothyronine (T3), thereby shaping hormone-dependent transcriptional responses to environmental and social context (Aksoy et al., 2022). In the medial amygdala, *Crym* overexpression increases transcriptional plasticity and modulates behavioral responses to social isolation stress (Walker et al., 2020), and in our prior work *Crym* was dramatically elevated in the amygdala of male mice transitioning social status (Milewski et al., 2025). Although in the current study neuronal *Crym* expression differed substantially by social status between all groups, the direction of this effect varied by sex (higher in subordinate rather than dominant females). We observed another discordant effect of social status in the relative quantities of *Trh* glutamatergic neurons, where dominant females and subordinate males both had significantly heightened proportions of this neuronal subtype. Given that *Trh* expression in hypothalamic neurons is negatively regulated by T3 (Harris et al., 2001; Nillni, 2010), the link between increased *Crym* transcription and enhanced stabilization of available T3 in this cell population offers a potential mechanistic explanation for the corresponding increase in *Trh* neurons we observed.

In the present study, the enrichment of multiple cholinergic-related subtypes (*Lhx8* and *Chat* GABAergic) in dominant individuals further suggests that dominance engages neuromodulatory systems that support social information processing and neuroendocrine output. *Chat* encodes the enzyme choline acetyltransferase responsible for the synthesis of acetylcholine within cholinergic neurons (Oda, 1999), whereas LHX8 is a key determinant of cholinergic neuron differentiation and promotes vesicular acetylcholine transporter expression and acetylcholine release (Mori et al., 2004; Tomioka et al., 2014). Converging evidence indicates that forebrain cholinergic signaling contributes to social memory and is sensitive to social stress (Mineur et al., 2013; Okada et al., 2021; Picciotto et al., 2012). Indeed, we have previously demonstrated that dominant CD-1 males have increased expression of cholinergic signaling genes throughout several amygdala nuclei (Milewski, Lee, Rada, et al., 2025; Milewski, Lee, Young, et al., 2025). In contrast to the enrichment of these inhibitory subtypes of neurons, subordinate animals and females had increased proportions of *Pmch* glutamatergic and *Sst* GABAergic neurons. PMCH neurons, which produce melanin-concentrating hormone (MCH), have well-established roles in motivated behavior, feeding, and sleep-wake state regulation, and higher representation of *Pmch*-tagged neuronal nuclei in subordinates in our study is consistent with rank-dependent differences in energy balance and rest-activity reported in dominant versus subordinate CD-1 mice. In parallel, *Sst* GABAergic neurons regulate local excitability and behavioral-state dependent circuit inhibition, providing a plausible mechanism through which social status could shape the balance between behavioral activation and recovery (B. R. Lee et al., 2023; Zingg et al., 2017). Notably, increased expression of *Sst* and the corresponding receptor gene *Sstr3* (somatostatin receptor subtype 3) as well as a dramatic tripling in size of hypothalamic *Sst*+ neurons has been documented in dominant male cichlid fish (Hofmann & Fernald, 2000; Trainor & Hofmann, 2007). These results suggest a potentially conserved somatostatin-dependent mechanism through which social rank sculpts hypothalamic circuitry to support the behavioral, metabolic, and reproductive demands of maintaining dominance.

### The transcriptomic neuropeptidome: Neuropeptide co-expression patterns vary by sex and social status

We next characterized neuropeptide transcript co-expression across mPOA neuronal nuclei using a candidate set of 127 neuropeptide genes. Of these, 52 were expressed in at least one neuron, and neuropeptide expression was common across the dataset: 80% of neuronal nuclei expressed at least one of the three most prevalent neuropeptide genes, and neurons expressed an average of 2.21 neuropeptide genes. Such widespread neuropeptide expression and co-expression is consistent with prior single-cell atlases of the hypothalamus that emphasize combinatorial neuropeptide signatures as a common feature of hypothalamic neuronal diversity (Chen et al., 2017; Herget & Ryu, 2015; Moffitt et al., 2018a; Romanov et al., 2017), and parallels recent single-nucleus profiling of the male cichlid preoptic area reporting that >99% of neuronal nuclei express at least one neuropeptide and that dominant neurons express an average of 6.45 neuropeptide genes compared to 4.26 in subordinates (Miller-Crews & Hofmann, 2025). The distribution of neuropeptide co-expression in our dataset differed across both sex and social status. Dominant females exhibited the lowest levels of neuropeptide co-expression (average neuropeptides per neuron = 1.35), whereas subordinate males showed the highest (average = 2.87). This pattern contrasts directly from the cichlid POA/hypothalamus dataset, where dominant males exhibit a more integrated transcriptomic neuropeptidome and higher overall network connectivity relative to subordinates (Miller-Crews & Hofmann, 2025). This discrepancy may reflect differences in the scope of the brain regions sampled, as the cichlid dataset encompasses a broader hypothalamic region, or the more dramatic phenotypic and reproductive state changes associated with dominance transitions in cichlids. Nevertheless, taken together, both findings highlight the importance of neuropeptide co-expression networks in regulating social status across species.

### Oxytocin, vasopressin, and secretogranin II transcript co-expression is widespread

Within this broader transcriptomic neuropeptidome, *Oxt*, *Avp*, and *Scg2* emerged as the dominant contributors to differences between groups. These were the three most broadly expressed neuropeptide genes in our neuronal dataset (*Oxt* in 71% of neurons, *Scg2* in 28%, and *Avp* in 22%), and 80% of all nuclei expressed at least one of these transcripts. Broad but low-level neuropeptide transcription is a common feature of peptidergic circuits, where stored and translationally regulated mRNAs are thought to permit rapid, activity-dependent peptide production rather than constitutive secretion (Burbach, 2011; Campbell et al., 2017; Romanov et al., 2017). In hypothalamic neurons, oxytocin and vasopressin transcripts are under tight post-transcriptional control, enabling cells to maintain a reserve of peptide mRNA that can be selectively translated in response to physiological or social stimuli (Burbach, 2011). Similar patterns of widespread, low-abundance neuropeptide expression have been observed in other single-cell studies of hypothalamic circuits, suggesting that such transcriptional priming represents a general mechanism supporting flexible neuromodulatory responses (Burbach, 2011; Campbell et al., 2017; Herget & Ryu, 2015; Romanov et al., 2017). The largest neuronal cluster (#0) in our sample, for which *Oxt* and *Avp* were top marker genes, was prominently enriched in subordinate animals. However, this effect of social status was ubiquitous across neuronal subtypes. Subordinates of both sexes exhibited the highest proportions of *Oxt+* neurons (91% in both subordinate groups), whereas dominant males and females contained substantially fewer *Oxt+* neurons (55% and 63%, respectively). *Oxt* expression levels among *Oxt+* neurons were also higher in subordinates. *Avp* showed a stronger male status effect. Nearly half of subordinate male neurons were *Avp+* compared to 12% of dominant male neurons. These prevalence shifts were accompanied by pronounced changes in co-expression structure among the three transcripts. Dominant females exhibited the greatest proportion of *Oxt+* neurons expressing *Oxt* alone, whereas subordinates, particularly subordinate males, showed the highest proportion of neurons co-expressing all three of *Oxt*/*Avp*/*Scg2*. Covariance-based network analyses identified strengthened *Oxt*-*Avp* coupling as the primary driver of dominant/subordinate dissimilarity in both sexes.

In subordinate animals, increased transcription of these neuropeptide genes in mPOA neurons is accompanied by a significant enrichment of genes involved in protein stabilization and localization to the cell membrane as well as positive regulation of intracellular protein transport. The functional consequences of this transcriptional shift are yet unknown, and here we specifically measured preprohormone mRNA expression in the nucleus rather than mature mRNA expression throughout the cell. Despite this limitation, we propose that increased *Oxt* and *Avp* expression in subordinates may reflect a state of continuous transcription in neuronal nuclei allowing rapid post-transcriptional processing and translation in response to environmental stimuli. Oxytocin is produced in preoptic/hypothalamic nuclei and oxytocin receptor signaling in the mPOA has been directly implicated in social behaviors, consistent with this region acting as a key node linking social cues to neuroendocrine output (H.-J. Lee et al., 2009; O’Connell & Hofmann, 2011). Vasopressin modulates fluid balance as well as social investigation, anxiety-related behaviors, and aggression (Albers, 2012; Rigney et al., 2022). Our findings in the mPOA contrast with canonical paraventricular and supraoptic nuclei, where distinct magnocellular populations produce either oxytocin or vasopressin peptides almost exclusively (Madrigal & Jurado, 2021; Rae et al., 2022). The degree of *Oxt*-*Avp* transcript co-expression we observe is therefore striking relative to the classical segregation of magnocellular oxytocin- and vasopressin-producing populations in PVN/SON (Otero-García et al., 2016; Soumier et al., 2022; Yuan et al., 2019), where within-cell co-expression is generally low under baseline conditions but can increase in specific physiological states (e.g., lactation or osmotic challenge) (Berkhout et al., 2024; Mezey & Kiss, 1991; Telleria-Diaz et al., 2001). Secretogranin II, encoded by the gene *Scg2*, is a secretory protein that critically regulates large dense core vesicle packaging and facilitates the relatively delayed, sustained extracellular release of their contents such as neuropeptides, compared to synaptic vesicle transmission (Leenders et al., 1999; Ludwig, 1998; Mansvelder & Kits, 2000; Martucci et al., 2023; Ohbuchi et al., 2015; Park & Kim, 2009). The prominence of *Scg2* coupling with *Oxt* and *Avp* expression suggests that status-dependent co-expression may reflect coordinated tuning of LDCV dynamics that influence oxytocin and vasopressin release (Canosa et al., 2011; Mitchell et al., 2020). In addition, *Scg2* has been established as a candidate for mediating the suppression of contextual fear ensemble neurons in the hippocampus during fear extinction and facilitates the reorganization of hippocampal inhibitory synaptic inputs (Yap et al., 2021; Zuniga et al., 2024). Given this and the dominance-associated increase in *Scg2*+ neurons in cluster #8, we theorize that *Scg2* may also contribute to shaping inhibitory mPOA circuitry in accordance with the behavioral, metabolic, and reproductive demands of maintaining dominance. Overall, our data suggest that the transcriptional machinery of neuropeptidergic neurons is profoundly responsive to social stimuli (Dantzer et al., 1987; Tobin et al., 2010) and that social status reorganizes neuropeptide expression in a coordinated shift between more specialized (dominant) and more flexible (subordinate) neuropeptide transcriptional states (Bergquist & Ludwig, 2008; Kirchner et al., 2023; Yue et al., 2008).

Across neuronal clusters and HypoMap-defined subtypes, the status effect on *Oxt* and *Avp* was largely consistent, with dominant individuals generally exhibiting lower transcript abundance than subordinates. *Caprin2* glutamatergic neurons were a notable exception to this pattern being the only subtype in which dominant individuals showed similar *Oxt* and *Avp* expression levels among expressing neurons as well as similar proportions of *Caprin2/Oxt*+ and *Caprin2/Avp*+ neurons relative to subordinates. *Caprin2* neurons generally co-expressed *Avp* at greater proportions than the other subtypes and contained the highest *Avp* expression for all groups. CAPRIN2 regulates vasopressin release in response to osmotic changes (Konopacka et al., 2015), suggesting that these neurons may be specialized for regulating neuropeptide output in response to physiological demands (Bárez-López et al., 2022; Loh et al., 2017). Recent single-nucleus profiling of the cichlid fish POA/hypothalamus also identified a *Caprin2*-marked *Avp*/*Oxt*-expressing population with PVN-like features (Miller-Crews & Hofmann, 2025). This suggests the possibility that *Caprin2*-associated peptidergic neurons represent a conserved, homeostasis-weighted population in which core peptide transcription is relatively insulated from social-state modulation. Although *Caprin2* neurons consisted of <5% of our sample, subordinate animals had consistently greater proportions of these neurons, indicating that the social environment may still shape *Caprin2* neuron transcriptional profiles without altering within-cell *Oxt/Avp* co-expression patterns. This dissociation is particularly intriguing given that dominance is accompanied by pronounced changes in fluid balance and excretory behaviors in CD-1 mice, including increased drinking and scent-marking via major urinary proteins (W. Lee et al., 2017; Milewski et al., 2022). As increased urination is also exhibited by dominant male cichlids to signal social status during agonistic encounters and courtship (Simões et al., 2015), *Caprin2* neurons may serve as a homeostatic peptide-regulatory population even as most other mPOA neuronal populations exhibit coordinated, status-associated neuropeptide co-expression states.

Social rank was also associated with shifts in the representation of steroid receptor-expressing neurons in the mPOA, including *Esr1*, *Ar*, *Nr3c1*, and *Nr3c2*, even though transcripts for the corresponding receptors of the most abundant neuropeptide genes (*Oxt*, *Avp*, *Scg2*) were not detected at appreciable levels in our dataset (<0.5% of neurons). Subordinate males exhibited an increased proportion of *Esr1*+ neurons, consistent with evidence that estrogen receptor α signaling in hypothalamic circuits suppresses aggression and promotes social inhibition (Hashikawa et al., 2017; D. Wei et al., 2023), and suggesting that subordinate states may be accompanied by enhanced recruitment of estrogen-sensitive neuronal populations that constrain competitive behavior. Differences in the proportion of *Ar*+ neurons likewise point to status-dependent modulation of androgen-sensitive circuitry supporting male-typical aggression and reproductive physiology (Carver et al., 2021; Guthman & Falkner, 2022). Rank also influenced glucocorticoid (*Nr3c1*) and mineralocorticoid (*Nr3c2*) receptor-expressing populations, suggesting that status may shift the abundance of stress hormone-responsive neuronal populations (Abbott et al., 2003; Sapolsky, 2005; Williamson, Franks, et al., 2016). Finally, although RRHO patterns in most receptor-expressing subpopulations largely mirrored the global neuronal signature, *Ar*+ and *Nr3c1*+ neurons also showed a discordant signal, with enrichment for gene sets that were highest in dominant males as well as subordinate females, indicating sex-dependent rank effects within these receptor-defined neurons.

### Conclusions

In the present study, we showed that social status is associated with coordinated shifts in both mPOA neuronal composition and neuronal gene expression in male and female CD-1 mice. At the whole-neuron level, dominant males and females shared a strongly concordant status-associated expression signature enriched for synaptic and plasticity-related functions, while females exhibited substantially greater differential expression by status than males. At the cellular level, rank-dependent differences were concentrated in specific clusters and HypoMap-defined subtypes, including a highly status-sensitive cluster characterized by the expression of *Crym*/*Rarb* as well as broader rank-associated shifts across *Pmch*/*Sst*-expressing subtypes and multiple inhibitory and cholinergic-related populations. Neuropeptide co-expression was widespread and strongly reorganized by sex and status, with dominant individuals showing reduced *Oxt*/*Avp* abundance and a reorganization of *Oxt*-*Avp*-*Scg2* co-expression toward more specialized states. *Caprin2* glutamatergic neurons were an exception to this pattern, being the only neuron subpopulation to show similar neuropeptide expression in dominant and subordinate animals. Overall, these results demonstrate the mPOA as a key brain region where social rank is translated into sex-dependent shifts in cellular composition and coordinated neuroendocrine transcriptional states.

## Data and Code Availability

Data and code used in this study are available at www.github.com/kmmahach/snRNAseq_status and https://doi.org/10.5281/zenodo.19685732

## Acknowledgements

We would like to thank the UT Austin Genomics Sequencing and Analysis Facility (GSAF) for sequencing support.

## Supplementary Material

### Supplemental Tables

Supplemental Table 1 – Neuron RRHO quadrant genes

Supplemental Table 2 – Excitatory/inhibitory neuron GLM results

Supplemental Table 3 – Neuron cluster GLM results

Supplemental Table 4 – HypoMap subtype GLM results

Supplemental Table 5 – List of neuron cluster marker gene results

Supplemental Table 6 – Common DEGs across clusters

Supplemental Table 7 – Neuron DEG limma/permutation results

Supplemental Table 8 – Neuropeptidergic neuron GLM results

Supplemental Table 9 – Neuropeptide co-expression distribution comparison

Supplemental Table 10 – Neuropeptide co-expression GLM results

Supplemental Table 11 – Neuroendocrine receptor neuron GLM results

Supplemental Table 12 – Neuroendocrine receptor RRHO quadrant genes

Supplemental Table 13 – GO results for neuroendocrine receptor RRHO quadrant genes

### Supplemental Figures

**Supplemental Figure 1.**
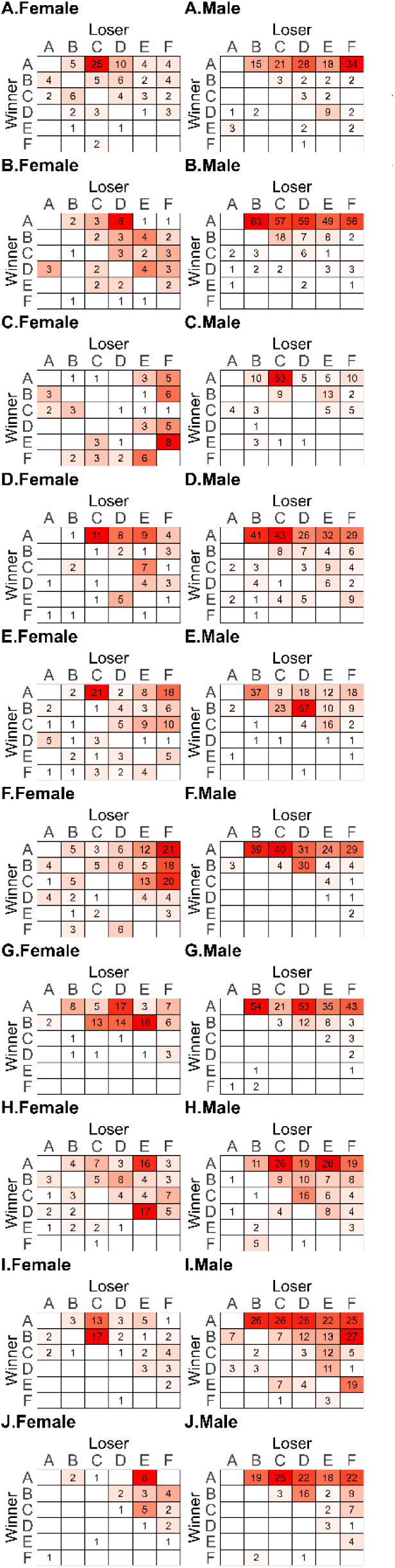
Frequency win-loss sociomatrices for all ten cohorts of each sex: female (A-J.Female) and male (A-J.Male). Total frequency of agonistic interactions between all pairs of individuals across all cohorts over the entire observation period.

**Supplemental Figure 2.**
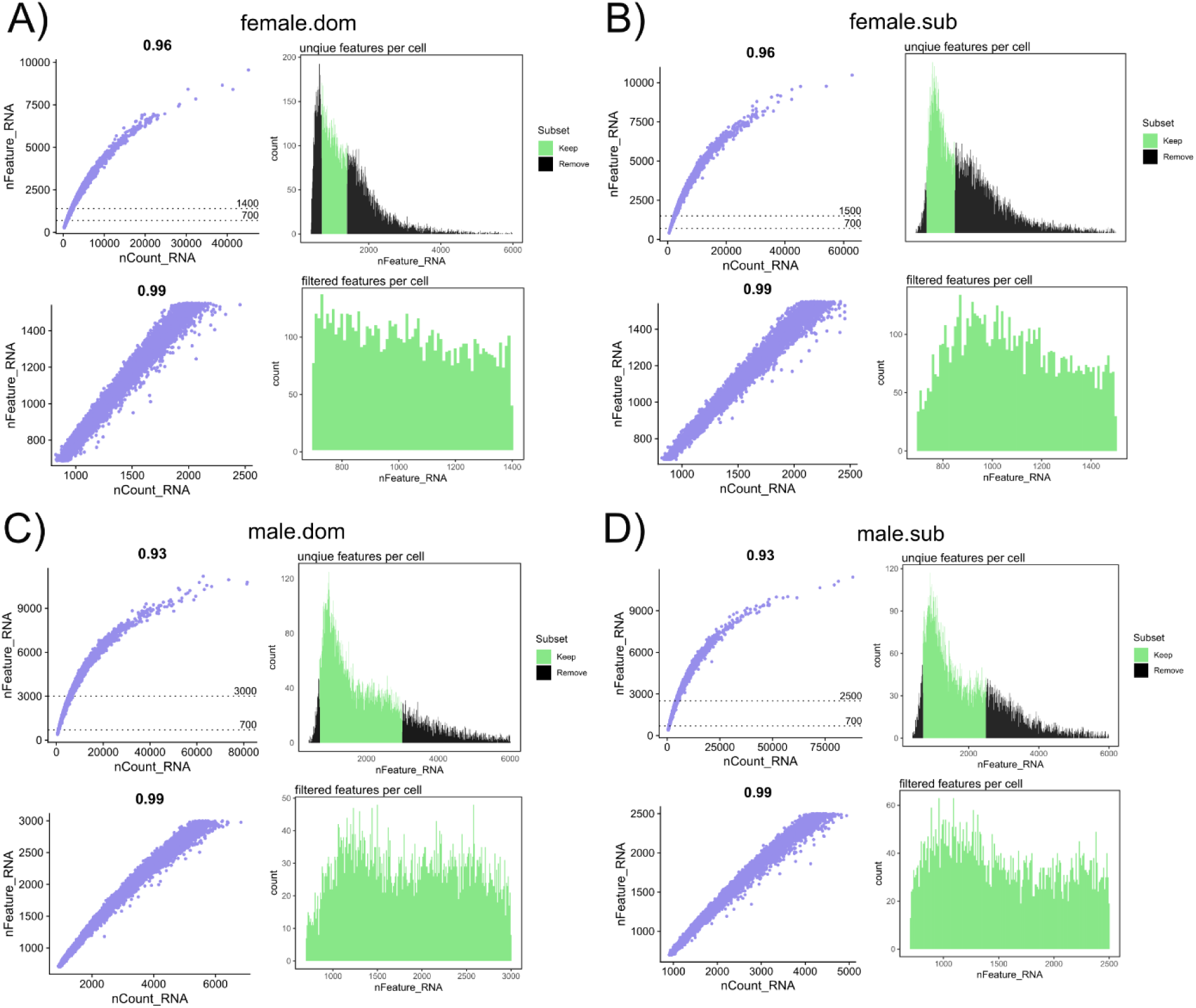
Quality control of CellRanger output for **A)** dominant females, **B)** subordinate females, **C)** dominant males, and **D)** subordinate males. Histograms (<u>right</u>) of the sample pool distribution including the targeted 10,000 nuclei (green), which were selected by excluding nuclei that expressed too many or too few unique genes (black). The selected nuclei demonstrated high positive correlation (<u>left</u>, r^2^ = 0.99) between the amount of Unique Molecular Identifiers (nCount_RNA) per nucleus and the number of distinct genes per nucleus (nFeature_RNA).

**Supplemental Figure 3.**
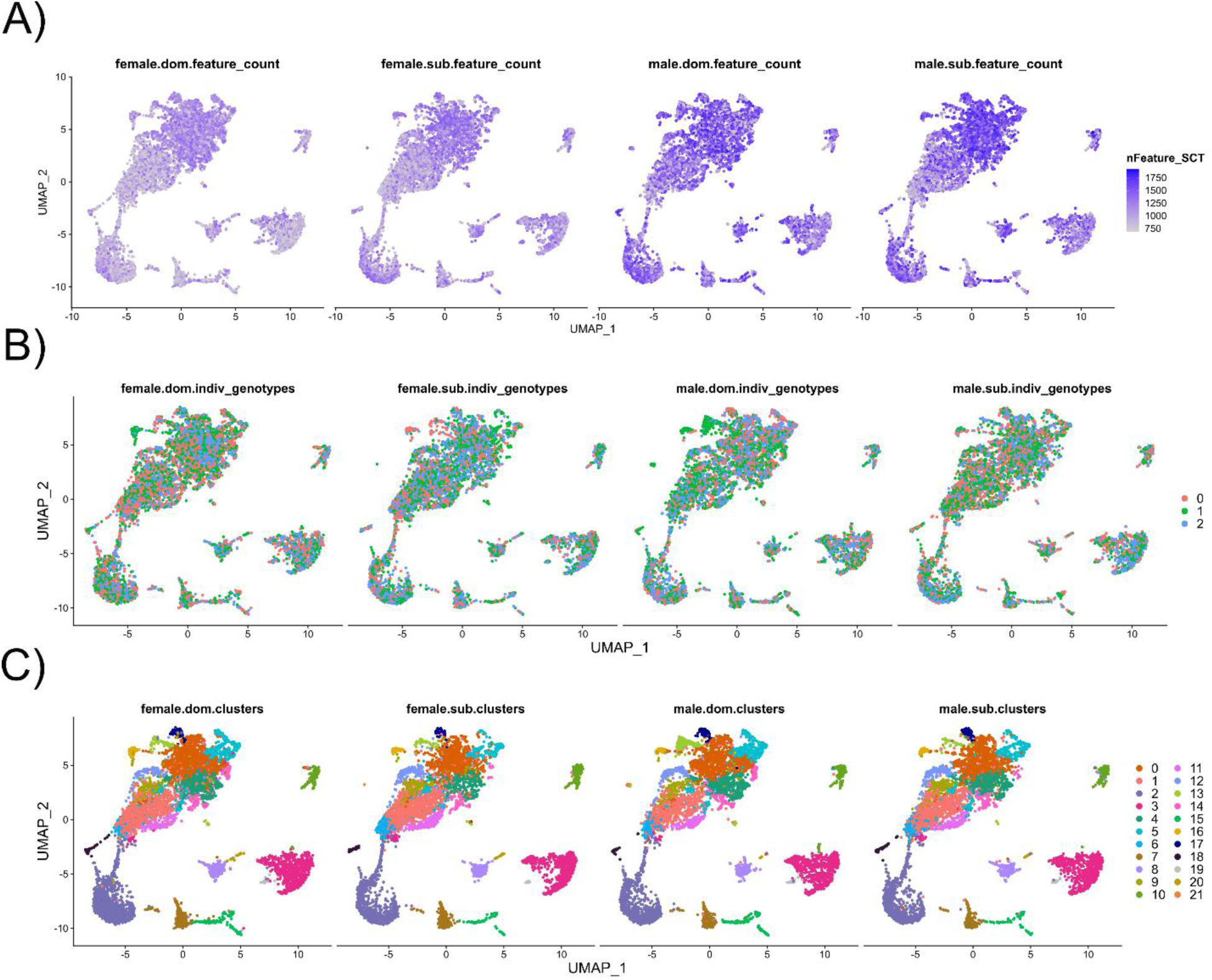
Quality control of normalization and integration across the four datasets (dominant females, subordinate females, dominant males, and subordinate males). **A)** UMAP plots with the distribution of feature counts per nucleus, where color intensity corresponds to the number of distinct genes (features). **B)** UMAP plots indicating the distribution of individuals among the three genotypes for each sample pool. Color indicates assignment to one of three putative genotypes within the sample pool. **C)** UMAP plots validating the successful integration of cell types across samples, with colors indicating statistically distinct cell clusters assigned post-integration.

**Supplemental Figure 4.**
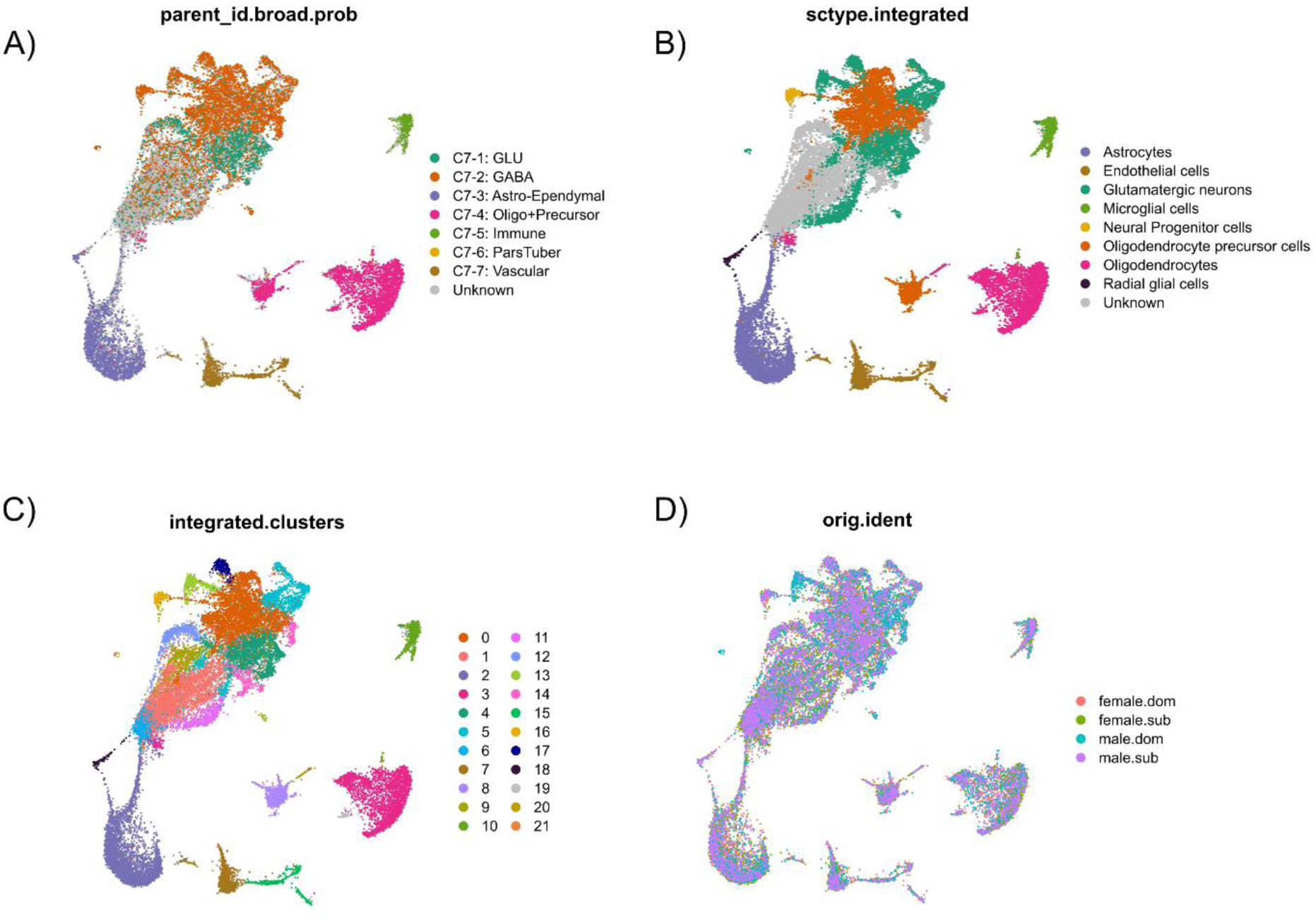
UMAP projections of all nuclei post-integration with broad cell type annotations predicted with either **A)** HypoMap or **B)** ScType. Nuclei with low confidence prediction scores (**≤** 0.75) were designated ‘unknown’. **C)** Distinct cell cluster assignments of all nuclei. **D)** Distribution of experimental groups by sample pool.

**Supplemental Figure 5.**
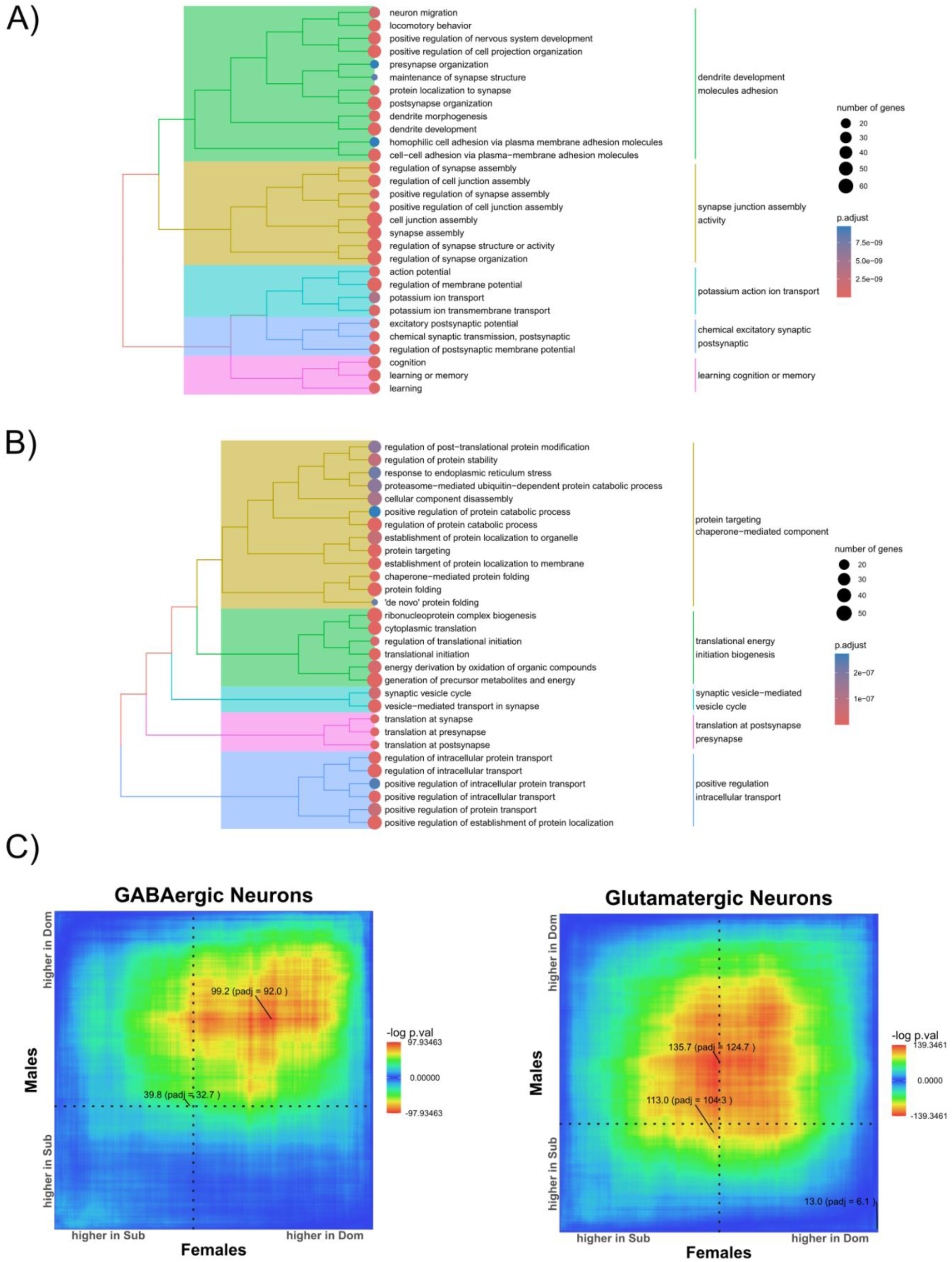
GO analysis results among genes enriched by social status across sex for either **A)** dominants or **B)** subordinates. Displayed are the top biological process (BP) categories and the hierarchical organization of semantic similarities between significant BP terms. **C)** RRHO plots of concordant and discordant gene expression patterns independently for inhibitory (<u>left</u>) and excitatory (<u>right</u>) neurons.

**Supplemental Figure 6.**
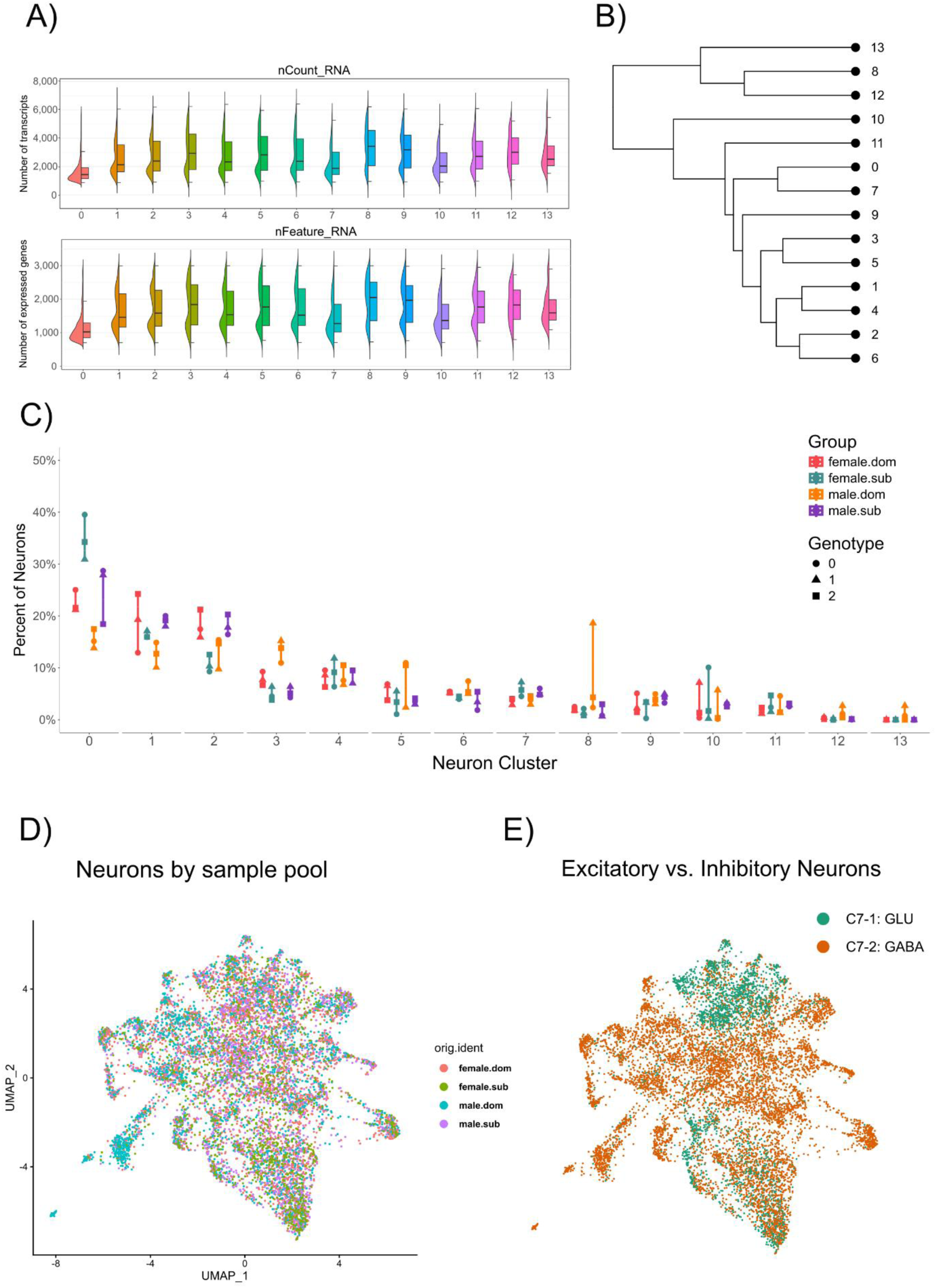
Sample composition and relationships between 14 neuron-specific clusters formed after excluding all other cell types. **A)** Quality control metrics for neuron clusters with distributions of total reads (<u>top</u>) and distinct genes (<u>bottom</u>) per nucleus. **B)** Hierarchical clustering of indicating similarity between neuron-specific clusters. Comparison of cluster composition across samples for **C)** percentage of nuclei. **E)** UMAP with sample pool of origin for all neuronal nuclei and **F)** corresponding excitatory/inhibitory neuron distribution.

**Supplemental Figure 7.**
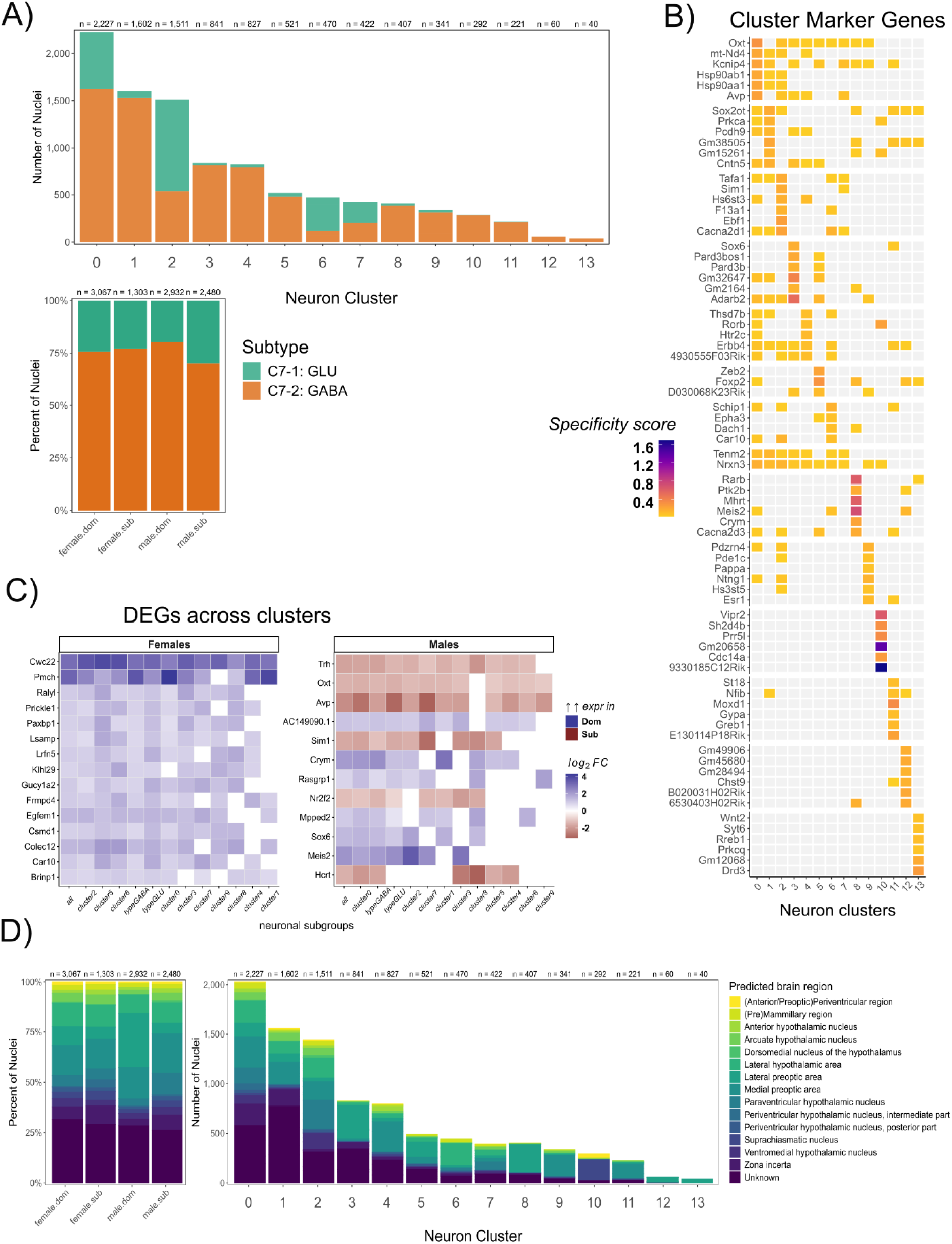
Neuron cluster attributes. **A)** Distribution of excitatory and inhibitory neurons across clusters. **B)** Heatmap of the top 6 marker genes (rows) per neuron cluster (columns) with color scaled by specificity score (*log_2_FC* * (pct1/pct2) * relative cluster size). Each marker gene is only represented once. **C)** Heatmap of genes for each sex that were differentially expressed by social status and their corresponding cluster-specific *log_2_FC* values (*eFDR* < 0.05). Genes (rows) are ordered (top to bottom, descending) by the number of clusters/subtypes in which they are differentially expressed; neuron subtypes (columns; including clusters and combined excitatory/inhibitory) are ordered (left to right, descending) by the number of common differentially expressed genes. Genes listed are the top 1% in each sex that were differentially expressed across the most clusters/subtypes, excluding those lacking samples from all individuals. **D)** Predicted regional origin of neurons by cluster.

**Supplemental Figure 8.**
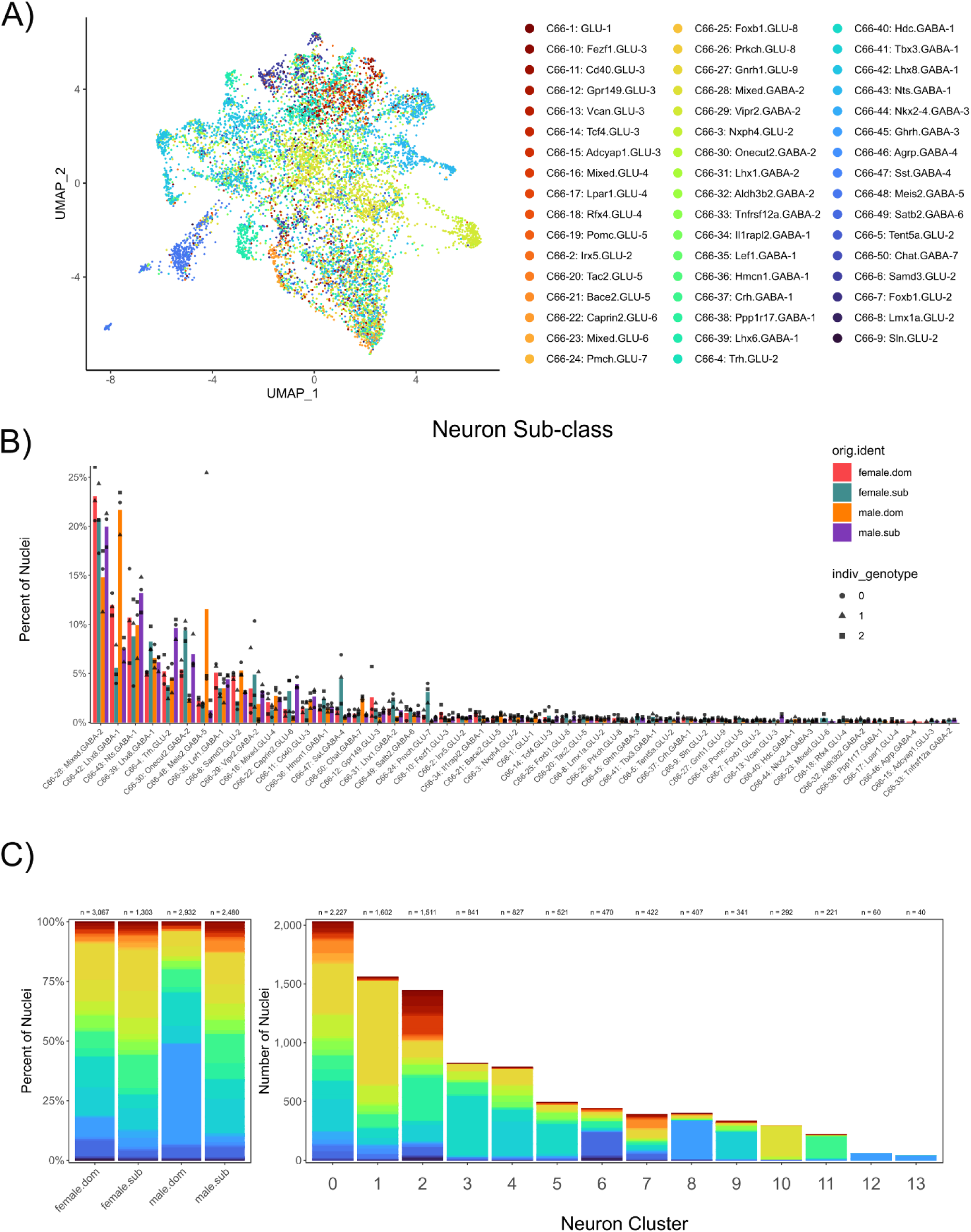
Neuron attributes and predicted sub-classifications. **A)** UMAP with distribution of predicted neuronal sub-classes. **B)** Bar plot with distribution of neurons across assigned sub-class colored by group with points for each individual in the sample pool. **C)** Percent of neuronal nuclei in each sub-class by group (left) and number of cells in each sub-class by neuron cluster (right). Colors match those denoted in the figure legend for *S8A*.

**Supplemental Figure 9.**
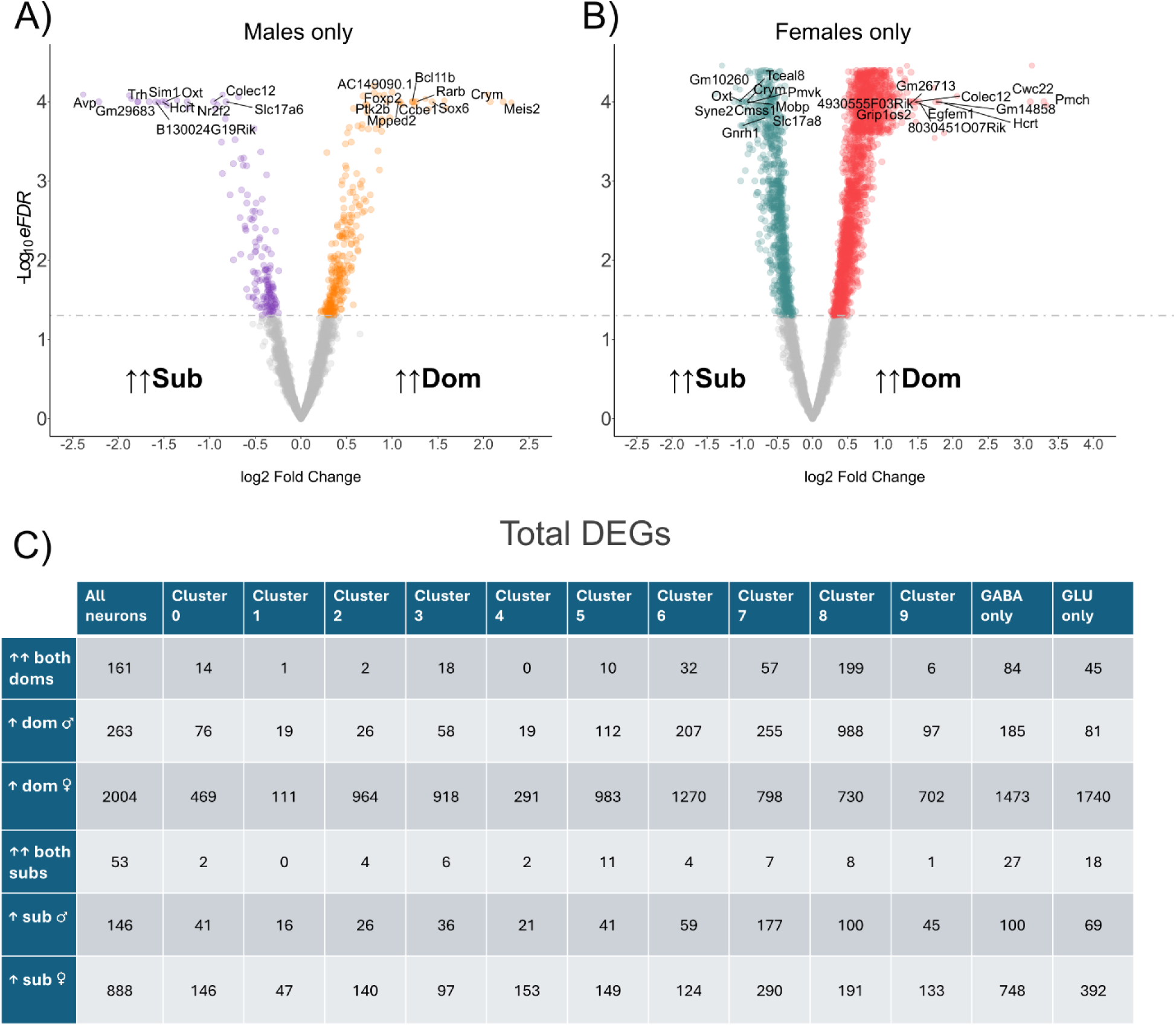
Genes differentially expressed by social status. **A)** Differentially expressed genes in male neurons only. **B)** Differentially expressed genes in female neurons only. Top 10 DEGs with largest *log_2_FC* values and smallest *eFDR* for either direction labeled on plot. Color indicates direction and significance. Gray genes did not meet our threshold for significance after permutation testing for differential expression with *limma* (*eFDR* < 0.05; permutations = 10,000). Points with -log_10_*eFDR* values > 3.5 are vertically jittered to prevent visual stratification of very low *eFDR* values, and *eFDR* values equal to zero (indicating that none of the 10,000 permutations produced a lower p-value than the true group assignments) are rounded up to 0.0001 for visualization. **C)** Total of DEGs in either direction (higher in dominants or subordinates) for males, females, and their intersection (*eFDR* < 0.05 & |*log_2_FC*| > 0.2).

**Supplemental Figure 10.**
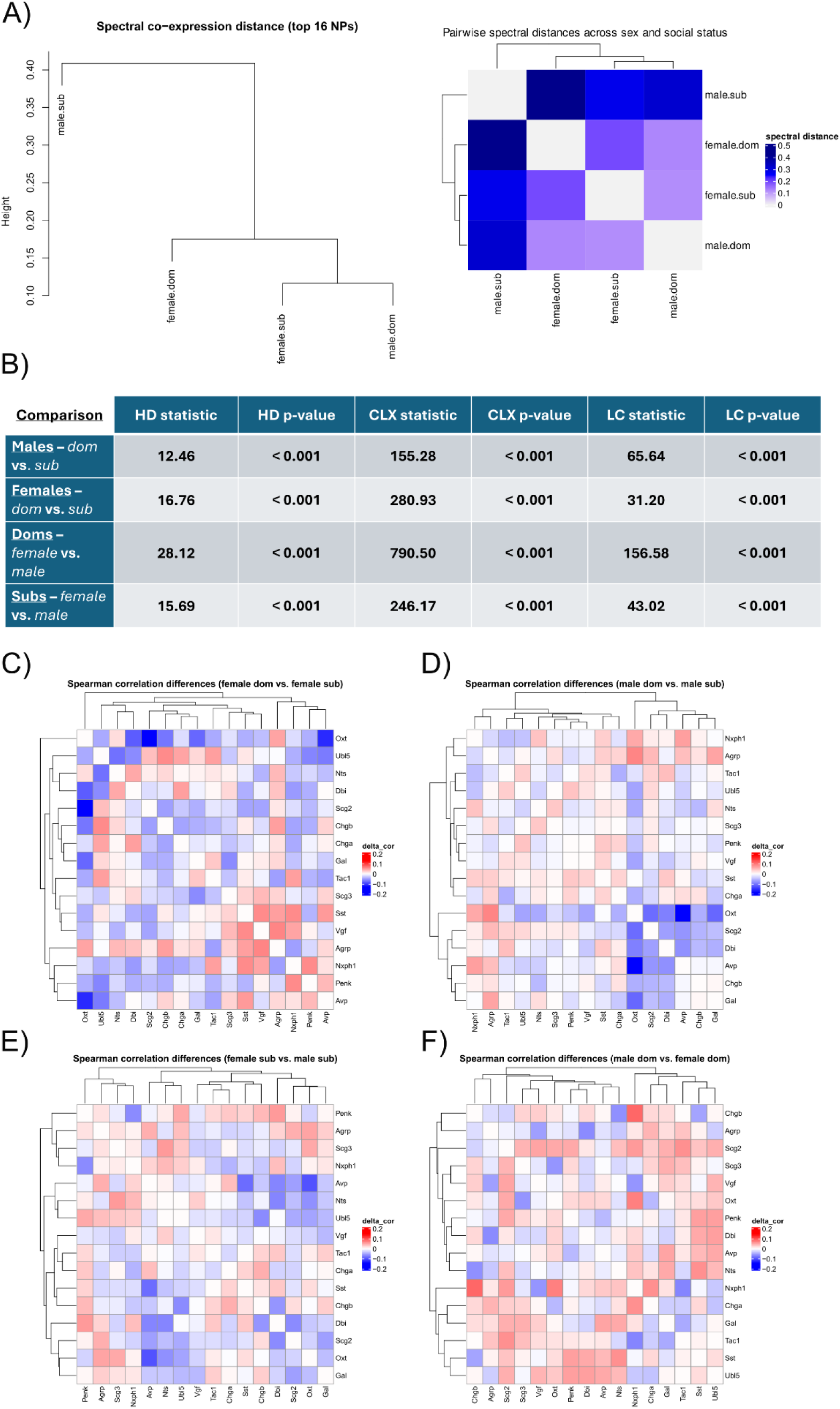
Comparison of neuropeptide co-expression similarity of the top 16 neuropeptides (expressed in >1% of neuronal nuclei). **A)** Hierarchical clustering was performed based on covariance matrix similarity across sample pools and visualized as a dendrogram of spectral distance (<u>left</u>) and as a heatmap (<u>right</u>) with all pairwise relationships derived from the co-expression patterns of top 16 neuropeptides. **B)** Three comparison statistics (HD, CLX, and LC) were used to establish global differences between covariance structures of all 52 expressed neuropeptide genes. **C)** Spearman correlation coefficients were calculated for each pair of neuropeptides from their normalized expression values, and the difference was taken between each pair of coefficients for the following group comparisons: Dominant females – subordinate females, **D)** Dominant males – subordinate males, **E)** Subordinate females – subordinate males, and **F)** Dominant males – dominant females.

**Supplemental Figure 11.**
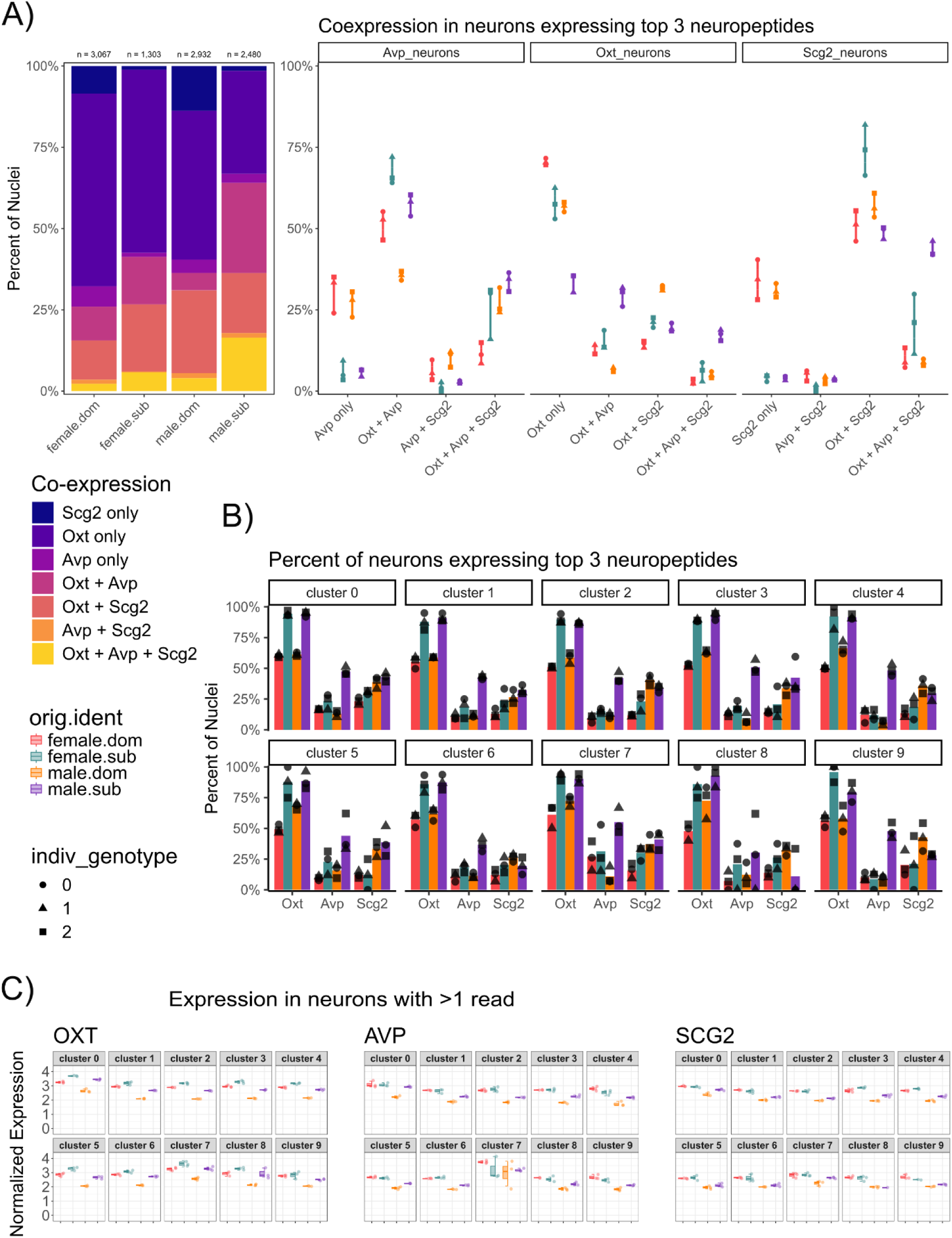
O*x*ytocin, *vasopressin*, and *secretogranin2* expression across clusters and neuronal subtypes. **A)** Percentages of neuronal nuclei in each sample pool that express any combination of *Oxt*, *Avp*, and *Scg2*. **B)** Percentages of neuronal nuclei that express each of these genes as a proportion of the neurons in each cluster. **C)** Normalized expression values of the top 3 neuropeptide genes only in neuronal nuclei that contain at least two raw reads (>1) of that transcript.

**Supplemental Figure 12.**
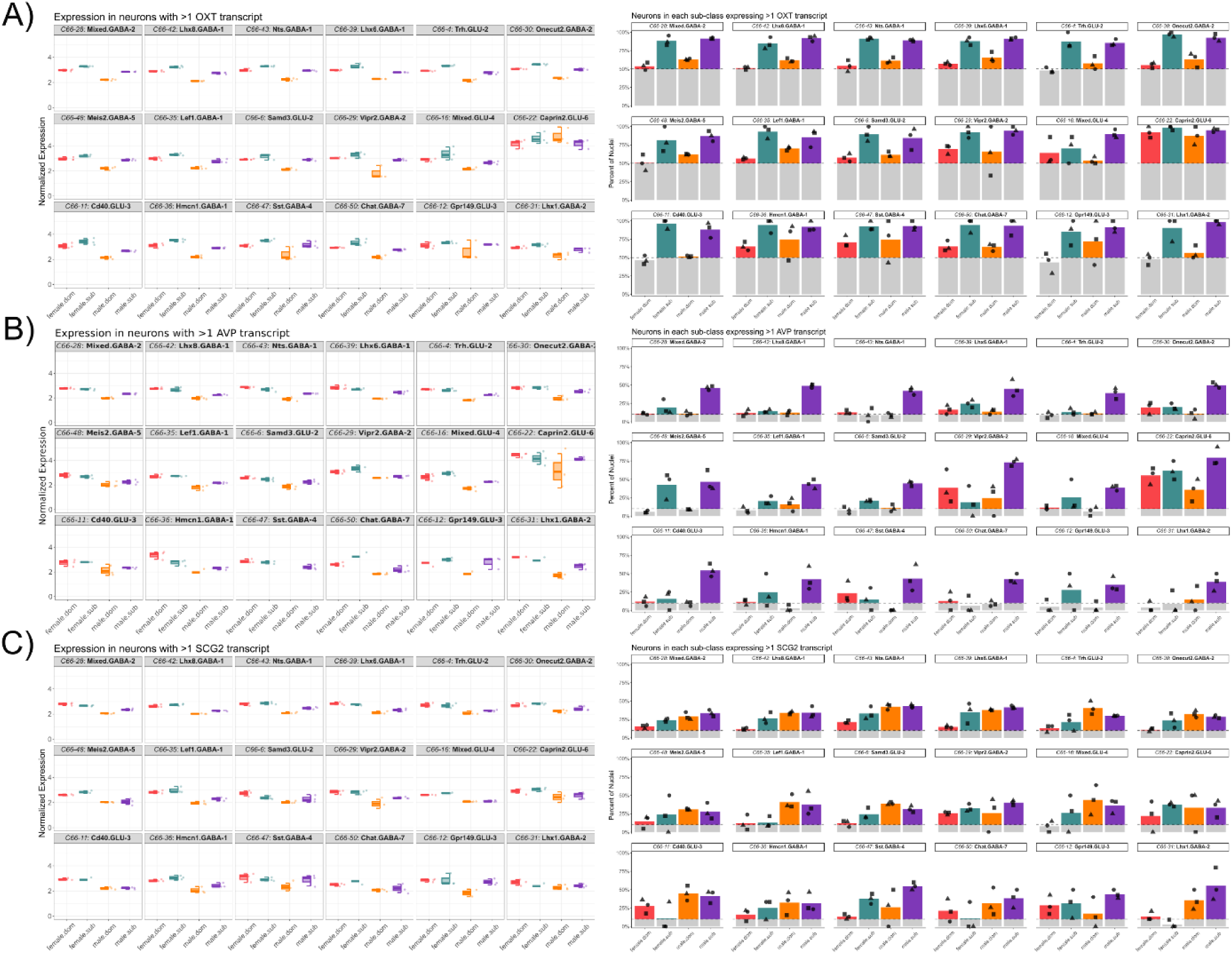
Normalized expression (<u>left</u>) of **A)** *Oxt* **B)** *Avp* and **C)** *Scg2* with the proportion of neurons (<u>right</u>) expressing >1 raw read in all neuronal sub-classes that comprise >1% of neurons (ordered as in *S8B*).

**Supplemental Figure 13.**
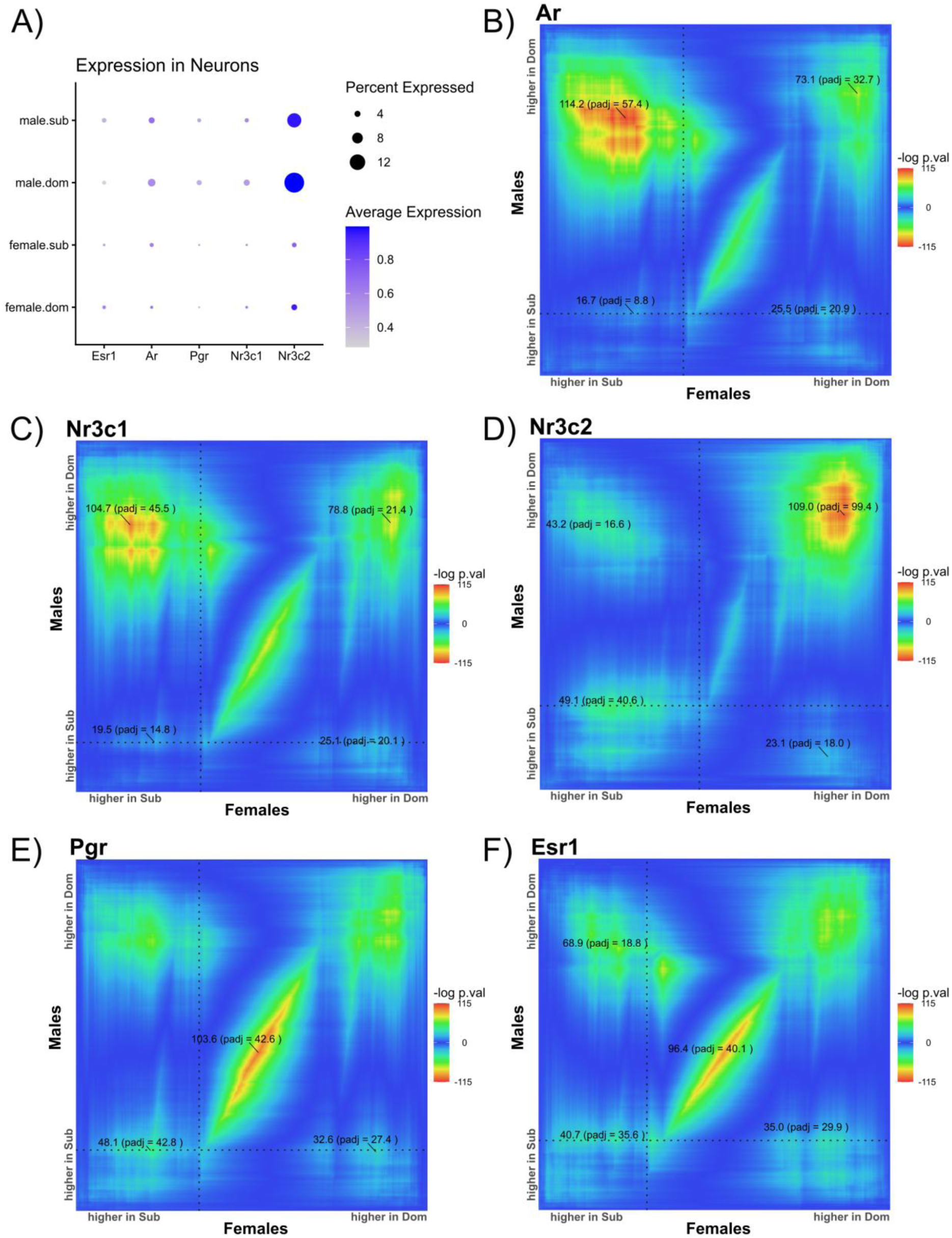
Hormone receptor expression in neurons. **A)** Dot plot; color indicates the average normalized expression values for these transcripts in all neurons for a sample pool and dot size indicates the percentage of neurons that express two or more (>1) reads of that gene. **B)** RRHO results comparing differential gene expression patterns across sex and between social status in nuclei expressing >1 transcript of the following steroid receptors: Androgen receptor (*Ar*), **C)** Glucocorticoid receptor (*Nr3c1*), **D)** Mineralocorticoid receptor (*Nr3c2*), and **E)** Progesterone receptor (*Pgr*), **F)** Estrogen receptor alpha (*Esr1*).

